# A microtubule RELION-based pipeline for cryo-EM image processing

**DOI:** 10.1101/673566

**Authors:** Alexander D. Cook, Szymon W. Manka, Su Wang, Carolyn A. Moores, Joseph Atherton

**Author notes:** Corresponding author Joseph Atherton.

## Abstract

Microtubules are polar filaments built from αβ-tubulin heterodimers that exhibit a range of architectures *in vitro* and *in vivo*. Tubulin heterodimers are arranged helically in the microtubule wall but many physiologically relevant architectures exhibit a break in helical symmetry known as the seam. Noisy 2D cryo-electron microscopy projection images of pseudo-helical microtubules therefore depict distinct but highly similar views owing to the high structural similarity of α- and β-tubulin. The determination of the αβ-tubulin register and seam location during image processing is essential for alignment accuracy that enables determination of biologically relevant structures. Here we present a pipeline designed for image processing and high-resolution reconstruction of cryo-electron microscopy microtubule datasets, based in the popular and user-friendly RELION image-processing package, **<u>Mi</u>crotubule <u>R</u>ELION-based <u>P</u>ipeline (MiRP)**. The pipeline uses a combination of supervised classification and prior knowledge about geometric lattice constraints in microtubules to accurately determine microtubule architecture and seam location. The presented method is fast and semi-automated, producing near-atomic resolution reconstructions with test datasets that contain a range of microtubule architectures and binding proteins.

**Abbreviations:** MiRP, Microtubule RELION-based Pipeline; cryo-EM, cryo-electron microscopy; MT, microtubule; CTF, contrast transfer function; PF, protofilament.

## Introduction

Structure determination using cryo-electron microscopy (cryo-EM) is a powerful and widely applicable methodology. Developments in both hardware and software have led to recent radical improvements in attainable resolutions, expansion of the types of samples that can be studied, and in experimental throughput (Kuhlbrandt, 2014). While a number of groups continue to develop and implement imaging processing methods, the accessibility of image processing software for non-expert users has been an important area of development. With intrinsic improvements in data quality, any image processing pipeline in principle requires fewer interventions and can incorporate robust automation of many steps; this has helped widen access to cryo-EM for structure determination. RELION (REgularised LIkelihood OptimisatioN) is a well-known and widely used software package, which implements a Bayesian approach to statistical modelling of cryo-EM data. It has been developed to allow new users to determine structures, while also allowing more sophisticated interventions by experienced practitioners (Scheres, 2012). It is open-source, actively maintained and updated, and has an engaged and knowledgeable community of users. It’s most recent release (v3.0) incorporated Bayesian polishing and per particle CTF correction (Zivanov et al., 2018).

Microtubules (MTs) are cytoskeleton polymers built from αβ-tubulin heterodimers that associate head-to-tail to form polar protofilaments (PFs) and laterally to form the hollow MT wall. They are central to many aspects of cell biology, acting as tracks for molecular motors and generating force via their dynamic growth and shrinkage. Structural studies have provided key insight into MT properties and functions and, given their size and complexity, EM has always been a vital tool in studying them (Manka and Moores, 2018a; Nogales and Zhang, 2016; Wade and Chretien, 1993) In particular, cryo-EM reconstructions of MTs in complex with a diverse array of binding partners have been important in understanding the distinct interaction modes of MTs with their binding partners and, in turn, in revealing mechanisms by which MTs are regulated.

The organisation of αβ-tubulin dimers within the MT wall is well-defined, although *in vitro* polymerised MTs often display a range of PF architectures which vary according to polymerisation condition and can be hard to differentiate visually (Pierson et al., 1978; Wade et al., 1990). Furthermore, the alternating α- and β-tubulin subunits are structurally very similar and are thus challenging to distinguish except at near-atomic resolutions. Binding partners attached every tubulin dimer act as fiducial markers and greatly facilitate structural discrimination between α- and β-tubulin (Zhang and Nogales, 2015).

Much of the early cryo-EM structural work on MTs focused on the relatively small subset of MTs with strict helical symmetry, in which the MT wall is entirely built of homotypic α-α and β-β contacts (Hirose et al., 1997; Kikkawa et al., 1995; Sosa et al., 1997). Fourier-based helical processing methods - which also depend on the long-range order of the MT polymer - were also typically employed. However, as is true for many biological polymers, MTs exhibit sufficient flexibility and distortion within the lattice that use of purely Fourier-based methods can limit the resolution of the final structure (Egelman, 2000). Furthermore, the structures of most MTs polymerised *in vitro*, as well as those found in many *in vivo* situations, include a single discontinuity - called the seam - in the otherwise helical arrangement of their subunits. This seam is composed of heterotypic α-β and β-α lateral contacts and standard Fourier helical methods cannot be used to determine the structure of such MTs.

To allow disorder within a single polymer to be accounted for, and to support structure determination of non-helical architectures, MTs in cryo-EM images are now more frequently treated as linear sets of “single particles”(Li et al., 2002). This allows each “single particle” piece of MT wall to be processed more or less independently, and for seam finding to be incorporated into the processing steps. In noisy, low electron dose 2D cryo-EM images of MTs, identification of the seam is a non-trivial computational task, given its overall similarity to the majority, homotypic lateral contacts in the MT wall. Nevertheless, several groups have successfully implemented MT “single particle” methods (Sindelar and Downing, 2007; Zhang and Nogales, 2015), thereby solving the structure of pseudo-helical MTs to near-atomic resolution (Kellogg et al., 2018; Manka and Moores, 2018b; Vemu et al., 2017; Vemu et al., 2016; Zhang et al., 2015; Zhang et al., 2018; Zhang et al., 2017), including in the absence of fiducial marker-like binding partners (Zhang et al., 2018).

Inspired by these previous studies and by the utility and popularity of RELION, we developed a <u>M</u>icrotubule <u>R</u>ELION-based <u>P</u>ipeline (MiRP). We describe here its organisation, quality-control outputs and application to number of different *in vitro* MT samples.

## Methods

### Protein expression and purification for cryo-EM

Recombinant human CAMSAP1 residues 1474-1613 encompassing the CKK domain (HsCKK) was purified from *E. coli* by the Akhmanova lab as described previously (Atherton et al., 2017a). tsA201 cell tubulin was purified from tsA201 cell cultures via tubulin TOG1 affinity and polymerisation cycling by the Roll-Mecak lab as described previously (Atherton et al., 2017a; Vemu et al., 2017; Vemu et al., 2014; Widlund et al., 2012). Recombinant mouse MKLP2 residues 25–520 including the motor domain and neck-linker was purified from *E. coli* by the Houdusse group as described previously (Atherton et al., 2017a).

Doublecortin (DCX) comprises two globular pseudo-repeats: NDC and CDC. Using a combination of standard PCR and restriction enzyme methods we generated a variant with two NDC repeats (NDC-NDC) in a pNic28-Bsa4 vector (Structural Genomics Consortium, Oxford, UK), which appends a TEV protease-cleavable His-tag on the N-terminus. The protein was expressed in BL21 Star (DE3) *E. coli* cells (Invitrogen). The cells were lysed by sonication in a lysis buffer (50 mM Na2HPO4 pH 7.2, 300 mM NaCl, 10 mM imidazole, 10% glycerol, 2 mM DTT) supplemented with protease inhibitor cocktail (cOmplete Cocktail Tablet, Roche/Sigma Aldrich). The lysate was clarified by centrifugation and passed through a HisTrap HP column (GE Healthcare). The protein was then eluted with 10-250 mM imidazole gradient and the His-tag was cleaved off using His-tagged TEV protease expressed in-house. Both the cleaved His-tag and the His-tagged protease were removed from protein solutions by passage over loose nickel beads (GE Healthcare). The tag-free NDC-NDC was further purified with HiTrap SP HP ion exchange column (GE Healthcare) equilibrated in BRB80 buffer (80 mM PIPES [piperazine-N,N′-bis(2-ethanesulfonic acid)] pH 6.8, 1 mM EGTA [ethylene glycol-bis(β-aminoethyl ether)-N,N,N’,N’-tetraacetic acid], 1 mM MgCl2, 1 mM DTT [dithiotreitol]) and eluted with NaCl gradient (15-300 mM). Final purification and desalting were done by gel filtration through Superdex 200 size exclusion column (GE Healthcare) equilibrated in BRB80 buffer.

### Sample preparation for Cryo-EM

HsCKK decorated MTs were prepared as described previously (Atherton et al., 2019). Briefly, tsA201 cell tubulin was polymerised in BRB80 (80 mM PIPES, 2 mM MgCl2, 1 mM EGTA, 1 mM DTT, pH 6.8) with 1 mM GTP at 37 °C then stabilised with 1 mM paclitaxel. At room temperature, stabilised MTs were then diluted 1/10 in BRB20 (20 mM PIPES, 2 mM MgCl2, 1 mM EGTA, 1 mM DTT, pH 6.8), adhered to holey carbon EM grids (C-Flat, Protochips Inc.) pre-glow discharged in air, and washed twice in 1 mg/ml HsCKK in BRB20. The EM grids were then blotted and vitrified using a Vitrobot (FEI/Thermo Fisher Scientific).

MKLP2 decorated MTs were prepared as described previously (Atherton et al., 2017b). Briefly, bovine tubulin (Cytoskeleton Inc.) was polymerised in MES polymerisation buffer (100 mM MES, 1 mM MgCl2, 1 mM EGTA, 1 mM DTT, pH 6.5) with 5 mM GTP at 37 °C then stabilised with 1mM paclitaxel. At room temperature, stabilised MTs were adhered to holey carbon EM grids (C-Flat, Protochips Inc.), pre-glow discharged in air, and washed once in 60μM MKLP2-MD pre-incubated in BRB20 containing 2mM of ADP +AlF4, before blotting and vitrification in a Vitrobot (FEI/Thermo Fisher Scientific).

For preparation of MTs decorated with the NDC domain of DCX, 5μM bovine tubulin (Cytoskeleton Inc.) was co-polymerised at 37 °C for 30 minutes with 3μM of NDC-NDC in BRB80 buffer with 1 mM GTP. The resultant NDC-NDC MT solution was applied to glow-discharged Lacey grids (Agar) and incubated for 20 sec at room temperature. Then the grids were briefly blotted and 50 μM NDC-NDC solution in BRB80 buffer was applied to boost MT decoration. The grids were then transferred to Vitrobot (FEI/Thermo Fisher Scientific) and incubated there for 1 min at 30 °C and 95 % humidity, before blotting and plunge freezing in liquid ethane.

### Cryo-EM Data collection

For all datasets, low dose movies were collected manually using SerialEM software (Mastronarde, 2005) on a FEI Tecnai G2 Polara operating at 300kV with a direct electron detector. Data collection details for each dataset can be found in Table 1.

**Table 1.** Dataset and data collection details. *K2 summit direct electron detector from Gatan Inc. CA, USA. ^†^DE20 direct electron detector from Direct Electron, San Diego, CA. ^‡^With quantum post-column energy-filter (Gatan Inc. CA, USA), operated in zero-loss imaging mode with a 20-eV energy-selecting slit. ^§^C-Flat 2/2-4C from Protochips Inc. ^ǁ^Lacey carbon grids from Agar Scientific.

| Decorator | Tubulin Ligands | Grid type | EM and energy filtration | Detector and mode | Pixel size (Å/pixel) | Defocus range | Dose (e-/Å<sup>2</sup>, weighted) | Total exposure time and frame number |
| --- | --- | --- | --- | --- | --- | --- | --- | --- |
| CKK domain of CAMSAP1 | α-tubulin: GTP<br>β-tubulin: GDP, Taxol | 2 μm Holey Carbon<sup>§</sup> | Polara (300kV<sup>‡</sup>) | K2 summit<sup>*</sup> Counting at 5e-/pixel/sec | 1.39 | 0.5-3.5μm | 42 dose weighted | 16 secs<br>64 frames |
| Motor domain of MKLP2 | α-tubulin: GTP<br>β-tubulin: GDP, Taxol | 2 μm Holey Carbon<sup>§</sup> | Polara (300kV) | DE20<sup>†</sup> (Direct Electron) Linear | 1.53 | 0.5-3.0μm | 50 dose weighted | 1.5 secs<br>22 frames |
| N-DC domain of DCX | α-tubulin: GTP<br>β-tubulin: GDP | Lacey Carbon | Polara (300kV<sup>‡</sup>) | K2 summit<sup>*</sup> Counting at 5e-/pixel/sec | 1.39 | 0.5-2.5μm | 24 unweighted | 9 secs<br>36 frames |

### Cryo-EM Data Processing Using MiRP

Unweighted and dose-weighted motion corrected sums were generated from low-dose movies using MotionCor2 (Zheng et al., 2017) with a patch size of 5. Full dose sums were used for contrast transfer function (CTF) determination in gCTF (Zhang, 2016), then dose-weighted sums were used in particle picking, processing and generation of the final reconstructions.

The details of the following MT processing protocol used for all datasets are described in Table 2. Briefly, start-end coordinates of MTs were manually picked in RELION, 4x binned particles extracted, then corresponding ‘segment average’ images generated (Fig. 2). 4x binned segment averages were subjected to supervised 3D classification with alignment to 15 Å low-pass filtered synthetic references of MTs of different PF architecture (generated in Chimera using known helical parameters, Table 3, (Sui and Downing, 2010). Particles from each MT were then assigned a modal consensus PF number class (Fig. 3a) and 13 PF MTs were taken for further processing. Psi and Tilt angles were set to the priors calculated in PF number classification, whilst Rot angles and translations were reset to 0.

**Table 2.** Details of the MiRP procedure. *If performing refinement with helical symmetry these options are set to ‘yes’. The appropriate helical parameters need to be optimised for each dataset. Example starting parameters that have worked well in our hands are; *13pf:* Number of asymmetrical units: 13 or 12 (if 13 or 12 binding proteins in a helical turn respectively). Intial twist (deg). rise (A): −27.67 9.46 Central Z length (%): 30 Twist search - Min,Max,Step (deg): −27 −28 0.1 Rise search - Min,Max,Step (A): 9.4 9.7 0.1 *14pf:* Number of asymmetrical units: 14 or 13 (if 14 or 13 binding proteins in a helical turn respectively). Intial twist (deg). rise (A): −25.71 8.81 Central Z length (%): 30 Twist search - Min,Max,Step (deg): −25.2 −26.2 0.1 Rise search - Min,Max,Step (A): 8.6 9 0.1 † If this option does not stop the refinement at iteration 1, manually terminate the refinement after the first iteration and use the iteration 1 output star files.

| Pipeline stage | Relion operation | Custom MT operations | Rough computing times | Details/parameters |
| --- | --- | --- | --- | --- |
| 1. Pre-processing | a) Manual picking |  | ~1 day/500 micrographs (CPU) | Input micrographs: Selected motion corrected and dose-weighted micrographs after CTF determination<br>Pick start-end coordinates helices? Yes |
|  | b) Particle extraction |  | ~10mins for 100,000 particles (CPU, 40 MPI processors) | Input coordinates: Manual-picked coordinates<br>Particle box size: ~600Å<br>Rescale particles? Yes<br>Re-scaled size (pixels): particle box size/4<br>Extract helical segments: yes<br>Tube Diameter: 400Å<br>Use bimodal angular priors? Yes<br>Coordinates are start end only? yes<br>Cut helical tubes into segments? Yes<br>Number of asymmetrical units: 1<br>Helical rise (Å): 82Å |
|  |  | c) Segment average generation | ~ 2 hours for 100,000 particles (CPU) | Script in C shell<br>Uses IMOD and Relion functions |
| 2. Pf number sorting | a) 3D classification |  | ~ 20 hours for 100,000 particles (either CPU, 60 MPI processors or GPU, 5 MPI processors with 3 threads) | Input images STAR file: 4 x binned segment averages star file<br>Reference map: Star file with paths to microtubule references<br>Reference mask: None<br>Reference map is on absolute greyscale? No<br>Initial low pass filter (Å): 15Å (if references not already low-pass filtered)<br>Symmetry: C1<br>Do CTF-correction? Yes<br>Has reference been CTF-corrected? Yes<br>Have the data been phase flipped: No<br>Ignore CTFs until first peak: No<br>Number of classes: equal to number of input references<br>Reg param T: 4<br>Number of iterations: 1<br>Use fast subsets (for large datasets): No<br>Mask diameter (Å): 580<br>Mask individual particles with zeros? Yes<br>Limit resolution to E-step to (Å): 12Å<br>Perform image alignment? Yes<br>Angular sampling interval: 1.8°<br>Offset search range (pix): ~65Å<br>Offset search step (pix) 1<br>Perform local angular searches? No<br>Do helical reconstruction? Yes<br>Tube diameter - inner, outer (Å): -1 400<br>Angular search range - tilt, psi (deg): 15 10<br>Keep tilt-prior fixed: No<br>Range factor of local averaging: -1<br>Apply helical symmetry: No<br>Do local searches of symmetry: No<br>Additional arguments: --dont_check_norm |
|  |  | b) Class Unification | <1 minute (CPU) | Script in Perl and commands in R |
|  |  | c) Class extraction, zero shifts and ROT and reset PSI and TILT angles to priors | <1 minute (CPU) | Script in C shell |
| 3. Global search | a) 3D auto-refine (1st) |  | ~20 mins for 30,000 particles (GPU, 5 MPI processors, 3 threads) | Input images STAR file: A final .star file generated above of a single PF number class<br>Reference map: 4 x binned decorated synthetic reference of corresponding PF number<br>Reference mask: None<br>Reference map is on absolute greyscale? No<br>Initial low pass filter (A): 15Å (if references not already low-pass filtered)<br>Symmetry: C1<br>Do CTF-correction? Yes<br>Has reference been CTF-corrected? Yes<br>Have the data been phase flipped: No<br>Ignore CTFs until first peak: No<br>Mask diameter (A): 580<br>Mask individual particles with zeros? Yes<br>Use solvent-flattened FSCs? Yes<br>Angular sampling interval: 1.8°<br>Offset search range (pix): ~40Å<br>Offset search step (pix) 1<br>Local searches from auto-sampling: 0.9°<br>Do helical reconstruction? Yes<br>Tube diameter - inner, outer (A): -1 400<br>Angular search range - tilt, psi (deg): 15 10<br>Keep tilt-prior fixed: No<br>Range factor of local averaging: 2<br>Apply helical symmetry: No<br>Do local searches of symmetry: No<br>Additional arguments: --dont_check_norm --ignore_helical_symmetry --iter 1<sup>†</sup> |
|  |  | b) Zero shifts and ROT, reset PSI and TILT angles to priors | <1 minute (CPU) | Script in C shell |
| 4. Unique Phi angle Assignment | a) 3D auto-refine (2<sup>nd</sup>) |  | ~10 mins for 30,000 particles (GPU, 5 MPI processors, 3 threads) | Input images STAR file: The .star file generated in the previous step.<br>Reference map: 4 x binned decorated synthetic reference of corresponding PF number<br>Reference mask: None<br>Reference map is on absolute greyscale? No<br>Initial low pass filter (A): 15Å (if references not already low-pass filtered)<br>Symmetry: C1<br>Do CTF-correction? Yes<br>Has reference been CTF-corrected? Yes<br>Have the data been phase flipped: No<br>Ignore CTFs until first peak: No<br>Mask diameter (A): 580<br>Mask individual particles with<br>zeros? Yes<br>Use solvent-flattened FSCs? Yes<br>Angular sampling interval: 0.9°<br>Offset search range (pix): ~45Å<br>Offset search step (pix) 0.5<br>Local searches from auto-sampling: 0.5°<br>Do helical reconstruction? Yes<br>Tube diameter - inner, outer (Å): -1 400<br>Angular search range - tilt, psi (deg): 3 3<br>Keep tilt-prior fixed: No<br>Range factor of local averaging: 2<br>Apply helical symmetry: No<br>Do local searches of symmetry: No<br>Additional arguments: --dont_check_norm --ignore_helical_symmetry --iter 1† |
|  |  | b) Phi Unification | <1 minute (CPU) | Script in Python |
|  |  | c) Zero shifts and reset PSI and TILT angles to priors | <1 minute (CPU) | Script in C shell |
|  | d) 3D auto-refine (3<sup>rd</sup>) |  | ~10 mins for 30,000 particles (GPU, 5 MPI processors, 3 threads) | Input images STAR file: The .star file generated in the previous step, but with links to 'raw' particles (not segment averages).<br>Reference map: 4 x binned decorated synthetic reference of corresponding PF number<br>Reference mask: None<br>Reference map is on absolute greyscale? No<br>Initial low pass filter (Å): 15Å (if references not already low-pass filtered)<br>Symmetry: C1<br>Do CTF-correction? Yes<br>Has reference been CTF-corrected? Yes<br>Have the data been phase flipped: No<br>Ignore CTFs until first peak: No<br>Mask diameter (Å): 580<br>Mask individual particles with zeros? Yes<br>Use solvent-flattened FSCs? Yes<br>Angular sampling interval: 0.9°<br>Offset search range (pix): ~45Å<br>Offset search step (pix) 0.25<br>Local searches from auto-sampling: 0.5°<br>Do helical reconstruction? Yes<br>Tube diameter - inner, outer (Å): -1 400<br>Angular search range - tilt, psi (deg): 3 3<br>Keep tilt-prior fixed: No<br>Range factor of local averaging: 2<br>Apply helical symmetry: No<br>Do local searches of symmetry: No<br>Additional arguments: --dont_check_norm --ignore_helical_symmetry --iter 1† --sigma_rot 3 |
| 5. X/Y Shift Smoothing |  | a) X/Y Shift Smoothing | <1 minute (CPU) | Script in Python |
|  | b) 3D auto-refine (4<sup>th</sup>) |  | ~5 mins for 30,000 particles (GPU, 5 MPI processors, 3 threads) | Input images STAR file: The .star file generated in the previous step ('raw' particles).<br>Reference map: 4 x binned decorated synthetic reference of<br>corresponding PF number<br>Reference mask: None<br>Reference map is on absolute greyscale? No<br>Initial low pass filter (A): 15Å (if references not already low-pass filtered)<br>Symmetry: C1<br>Do CTF-correction? Yes<br>Has reference been CTF-corrected? Yes<br>Have the data been phase flipped: No<br>Ignore CTFs until first peak: No<br>Mask diameter (A): 580<br>Mask individual particles with zeros? Yes<br>Use solvent-flattened FSCs? Yes<br>Angular sampling interval: 0.9°<br>Offset search range (pix): ~35Å<br>Offset search step (pix) 0.5<br>Local searches from auto-sampling: 0.5°<br>Do helical reconstruction? Yes<br>Tube diameter - inner, outer (A): -1 400<br>Angular search range - tilt, psi (deg): 2 2<br>Keep tilt-prior fixed: No<br>Range factor of local averaging: 2<br>Apply helical symmetry: No<br>Do local searches of symmetry: No<br>Additional arguments: --dont_check_norm --ignore_helical_symmetry --iter 1<sup>†</sup> --sigma_rot 3 |
| 6. Refined segment averages | a) Particle extraction |  | ~4mins for 30,000 particles (CPU, 40 MPI processors) | Input Coordinates:<br>OR re-extract refined particles? yes<br>Refined particles star file: The .star file generated in the previous step.<br>Reset the refined offsets to 0? No<br>Re-center refined coordinates: Yes<br>Recenter on- X,Y,Z (pix): 0,0,0<br>Particle box size (pix) = same as previous extraction<br>Rescale particles? Yes<br>Re-scaled size (pixels): same as previous extraction<br>Extract helical segments = yes<br>Tube Diameter (A) = 400Å<br>Use bimodal angular priors? yes<br>Coordinates are start end only? No<br>Cut helical tubes into segments? Yes<br>Number of asymmetrical units: 1<br>Helical rise (A): 82Å |
|  |  | b) Refined segment average generation | <1 minute (CPU) | Script in C shell<br>Uses IMOD and Relion functions |
| 7. Seam Finding | a) 3D classification |  | ~10mins for 30,000 particles (CPU, 1 MPI processor, 9 threads) | Input images STAR file: 4 x binned refined segment averages star file<br>Reference map: Star file with paths to seam position and $\alpha/\beta$ register references<br>Reference mask: None<br>Reference map is on absolute greyscale? No<br>Initial low pass filter (A): 15Å (if references not already low-pass filtered)<br>Symmetry: C1<br>Do CTF-correction? Yes<br>Has reference been CTF-corrected? Yes<br>Have the data been phase flipped: No<br>Ignore CTFs until first peak: No<br>Number of classes: equal to number of input references<br>Reg param T: 4<br>Number of iterations: 1<br>Use fast subsets (for large datasets): No<br>Mask diameter (A): 580<br>Mask individual particles with zeros? Yes<br>Limit resolution to E-step to (A): 12A<br>Perform image alignment? No<br>Do helical reconstruction? Yes<br>Tube diameter - inner, outer (A): -1 400<br>Angular search range - tilt, psi (deg): 15 10<br>Keep tilt-prior fixed: No<br>Range factor of local averaging: -1<br>Apply helical symmetry: No<br>Do local searches of symmetry: No<br>Additional arguments: --dont_check_norm |
|  |  | b) Class Unification | <1 minute (CPU) | Script in Perl and commands in R |
|  |  | c) Class extraction | <1 minute (CPU) | Script in C shell |
|  |  | d) Phi/XY Correction | <1 minute (CPU) | Script in C shell |
| 8. High-resolution reconstruction | a) Particle extraction |  | ~10mins for 30,000 particles (CPU, 40 MPI processors) | Input Coordinates:<br>OR re-extract refined particles? yes<br>Refined particles star file: The .star file generated in the previous step. Reset the refined offsets to 0? No<br>Re-center refined coordinates: Yes<br>Recenter on- X,Y,Z (pix): 0,0,0<br>Particle box size (pix) = same as previous extraction<br>Rescale particles? No<br>Extract helical segments = yes<br>Tube Diameter (A) = 400Å<br>Use bimodal angular priors? yes<br>Coordinates are start end only? No<br>Cut helical tubes into segments? Yes<br>Number of asymmetrical units: 1<br>Helical rise (A): 82Å |
|  | b) 3D auto-refine (5<sup>th</sup>) |  | ~ 6 hours for 30,000 particles (either CPU, 60 MPI processors or GPU, 5 MPI processors with 3 threads) | Input images STAR file: The .star file generated in the previous step.<br>Reference map: 4 x binned decorated synthetic reference of corresponding PF number<br>Reference mask: None<br>Reference map is on absolute greyscale? No<br>Initial low pass filter (A): 15Å (if references not already low-pass filtered)<br>Symmetry: C1<br>Do CTF-correction? Yes<br>Has reference been CTF-corrected? Yes<br>Have the data been phase flipped: No<br>Ignore CTFs until first peak: No<br>Mask diameter (A): 460<br>Mask individual particles with zeros? Yes<br>Use solvent-flattened FSCs? Yes |
|  |  |  |  | Angular sampling interval: 0.9°<br>Offset search range (pix): ~6Å<br>Offset search step (pix) 0.5<br>Local searches from auto-sampling:<br>0.9°<br>Do helical reconstruction? Yes<br>Tube diameter - inner, outer (Å):<br>120 400<br>Angular search range - tilt, psi<br>(deg): 2 2<br>Keep tilt-prior fixed: No<br>Range factor of local averaging: -1<br>Apply helical symmetry: No*<br>Do local searches of symmetry:<br>No*<br>Additional arguments: --<br>dont_check_norm --sigma_rot 2 |

**Table 3.** Helical parameters used for protofilament number references.

| Protofilament Number | Rise (Å) | Twist (°) |
| --- | --- | --- |
| 11-3 | 11.1 | -32.5 |
| 12-3 | 10.2 | -29.9 |
| 13-3 | 9.4 | -27.7 |
| 14-3 | 8.7 | -25.8 |
| 15-4 | 10.8 | -23.8 |
| 16-4 | 10.2 | -22.4 |

4x binned segment averages of the single PF number class were then roughly aligned for a single iteration to a simulated 3D reference of an MT with appropriate PF number with decorating protein, filtered to 15 Å. Psi and Tilt angles were once again set to their corresponding newly assigned priors whilst Rot angles and translations were reset to 0. A second single iteration refinement was then performed to the same reference, with finer sampling of all angles, and local sampling of the Psi/Tilt angle.

For a given MT, the most commonly observed Rot angle was determined and assigned to all particles in the MT (Fig. 4a,b). To do this, a distance matrix of the Rot angles assigned to all particles was calculated, and clusters of angles within 8 ° of each other were extracted. 8 ° was used to ensure only Rot angles aligning the same particle-reference protofilament register were clustered. Linear regression was then performed on the most populated cluster, with the resulting slope and intercept used to calculate and impose the Rot angle for all particles. A third single iteration refinement was then performed to the same reference, with local sampling of the Rot/Tilt/Psi angle.

Next, X/Y shifts were smoothened to remove intermittent mis-translations along the MT axis (Fig. 4c,d). To do this, clusters of particles separated by mis-translations were created. Linear regression was performed on the most populated cluster, with the resulting slope and intercept used to calculate the X/Y shifts for all particles. A fourth single iteration refinement of raw particles was then performed to the same reference, with local sampling of all angles, and the X/Y shifts. Based on these alignment parameters, centred 4xbin particles were then re-extracted and new segment averages created.

To check and correct the MT-Rot angle allocation for each MT, a supervised 3D classification without alignment of centred segment averages was performed (Fig. 5). This was done using 15 Å low-pass filtered synthetic references of decorating protein density alone (excluding density corresponding to tubulin), rotated around the helical axis to represent all possible seam positions with or without a 41Å shift along the helical axis (Fig. 5a,b, 26 references total for a 13PF MT). Class allocations for particles within each MT were then set to a consensus corresponding to the most common class. Based on this consensus class allocation, the angles and corresponding translations along the helical axis were then adjusted for each MT particle.

At this stage, poor quality MTs were removed based on the internal consistency of their alignment parameters (see Fig. 6a). Unbinned raw particles were then extracted, centred by the above-determined translations and a local C1 auto-refinement was performed with restrained translations and psi/tilt/ranges. A 3D classification without alignment was performed to the resulting reconstruction (see Fig. 6b), and good classes were subjected to another local C1 auto-refinement performed with restrained translations and Rot/Tilt/Psi ranges. A round of Bayesian polishing was then performed, followed by another identical round of auto-refinement. CTF refinement was then performed, again followed by a further restrained round of auto-refinement with or without appropriate helical symmetry application and symmetry refinement. Symmetrised reconstructions were processed by B-factor application to local resolution cut-offs (global B-factor and local-resolutions were determined in RELION) and the central asymmetric unit opposite the seam was analysed (Fig. 7).

### Data availability

All the scripts for use with MiRP are publically available at http://GitHub.com. MiRP-derived cryo-EM density for full C1 reconstructions and symmetrised asymmetric units will be made available upon publication at the Electron Microscopy Data Bank (EMDB) under the following accession codes: 13PF CKK-MT, EMD-4643, 13PF MKLP2-MT, EMD-XXXX and 13PF NDC-MT, EMD-XXXX.

The 13PF MKLP2-MT model displayed in MiRP-derived density (Fig. 7) has been previously published (Atherton et al., 2017b) and deposited in the protein data bank (PDB) under the accession code 5ND4. The 13PF CKK-MT model was calculated using MiRP derived density and is described in a separate manuscript (Atherton et al., 2019). 13PF CKK-MT and 13PF NDC-MT models will be made available upon publication at the PDB under the following accession codes: 13PF CKK-MT, 6QUS and 13PF NDC-MT, XXXX.

## Results

### Overview of MiRP design

MTs polymerised *in vitro* are a heterogeneous population, typically composed of MTs of varying PF number architecture. For filamentous objects such as MTs, the three Euler angles refer to specific structural parameters: the Rot angle describes the rotation of the particle around the z-axis of the 3D reference, the Tilt angle describes the out-of-plane tilt, and the Psi angle the in-plane-rotation. The Rot, Tilt and Psi angles are also commonly termed φ, θ, and ψ, but here we use the former, RELION nomenclature. X and Y translations describe the image shifts necessary in the two in-plane dimensions required to align the particle to the reference. The objective of MiRP (Fig. 1) is to correctly identify the MT PF number architecture, Euler angles and translations for images of MTs. As will be described in more detail, the biggest challenges are identifying the Rot angle and translations, thus they are the focus of MiRP design.

**Figure 1.**
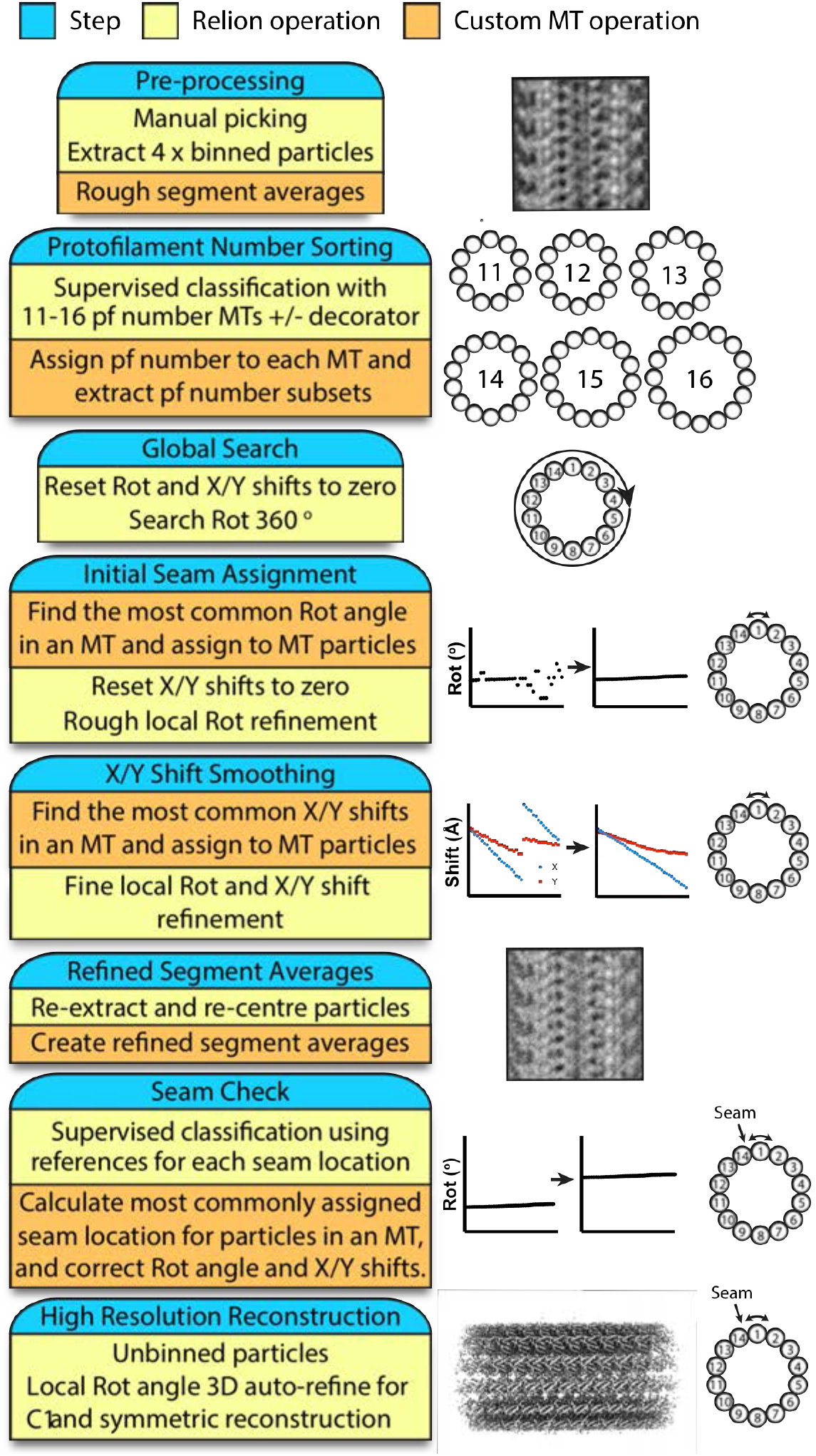
The MT Image processing RELION-based Pipeline (MiRP). Each step is marked in blue, in the same box as short summaries of the RELION operations (yellow), and custom MT operations (orange) involved in that step, described in more detail in the text.

In MiRP (Fig. 1), we use standard RELION operations that can be run from its graphical user interface. The procedure broadly follows classical steps for single particle analysis, starting with particle picking and extraction, then multiple rounds of particle classification and refinement, and a final high-resolution refinement from cleaned/optimised data. Each step however, has a specific objective designed expressly to deal with MT architecture heterogeneity and pseudo-symmetry, and is generally followed by custom operations that perform data analysis and/or manipulate the Euler angles, X/Y translations, or class assignment associated with that step (Fig. 1). The known structural constraints of MT polymers imply all neighbouring particles within any given MT will have the same architecture and related Euler angles and X/Y shifts. Therefore, angular and translational searches in RELION are restrained accordingly and custom operations, performed mostly using C shell and Python scripts, implement analyses and corrections based on this prior structural knowledge of the MT polymer.

Direct application of standard helical processing in RELION is not suitable for seam-containing MTs, first, because of the underlying assumption that there are multiple correct alignment solutions determined by the helical symmetry, rather than a single solution appropriate for a pseudo-symmetrical filament with a seam. Second, neighbouring particles within MTs are treated independently and thus useful constraints on class, Euler angle and translational allocations are not imposed, thereby failing to capitalise on prior structural knowledge of MTs.

The use of RELION’s graphical user interface makes it simple to perform and track MiRP steps and assess their success. It also allows easy access to useful features, such as per-particle CTF refinement, local resolution filtering, and post-processing procedures. Manual interventions are required at most steps, and therefore the procedure is not completely automated. To evaluate its efficacy and the importance of the manual interventions, we applied MiRP to three previously published exemplar MT datasets varying in their sample preparation and decorating protein (Table 1).

### Pre-processing

RELION’s manual picker can be used in helical mode to pick individual MTs (Fig. 2a). Only straight individual MT regions should be picked, making sure the centre of extracted boxes correspond to the centre of the MT, and avoiding curved, contaminated, or distorted regions (Fig. 2a) as well as overlapping MTs in bundles or at cross-overs. Distorted MT regions can occur for a variety of reasons including defects during tubulin polymerisation/depolymerisation, MT overlaps or contacts and ice thinning during the blotting and vitrification process (e.g. blue arrows in Fig. 2a, and (Atherton et al., 2018). MiRP is designed to utilise information from all neighbouring particles within an individual MT, and therefore performs optimally when the dataset is composed of long unbroken MTs without distortions or contamination. We have not been able to use RELION’s automated filament picking to both include all the best MT lengths while excluding non-ideal regions; therefore, manual picking is currently recommended. Whilst coordinates from other programs can be imported into RELION, we find RELION’s manual helical picker preferable because of its inbuilt low-pass filtering options and its filament picking straight-line traces that can be used to assess MT curvature.

**Figure 2.**
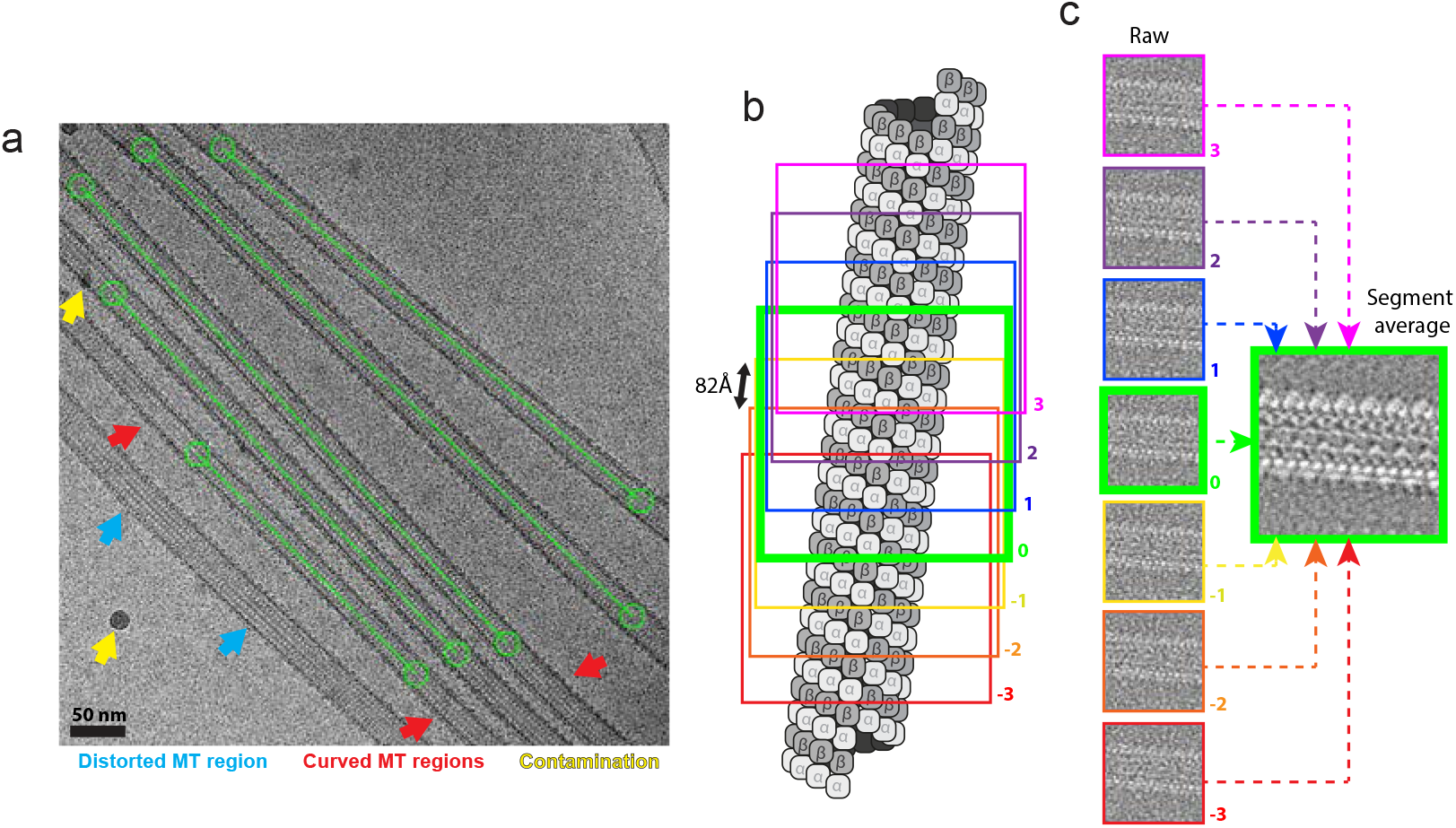
Pre-processing; manual picking and particle extraction strategy. a) “Helical” manual picking strategy is shown in an example micrograph from the CKK-MT dataset displayed using the RELION manual-picking GUI window. Start-end coordinates are selected (green circles) delineating desired MT lengths (connecting green lines) for extraction. Distorted or curved MT lengths (shown with blue and red arrows respectively) as well as contaminated areas (yellow arrows) are excluded. b) Particle extraction strategy illustrated on an MT diagram. Box size is set at roughly 600Å with a box separation of 82Å representing the dimer repeat distance along the helical axis. 7 example boxes are shown, where the central box (bold green) serves as the central particle for segment average generation. c) Segment average generation strategy. 3 adjacent particles either side of a central particle along the helical axis are averaged with a central particle to create a new central particle with a higher signal-to-noise ratio.

Particles are extracted such that each adjacent particle along a MT contains a unique asymmetric unit (tubulin dimer). However, this asymmetric unit is extracted in the context of a whole MT segment to allow accurate alignment. In practice, we extract particles with an inter-box distance of 82Å, representing roughly the dimer repeat distance, within a box size of around ~2x the MT diameter and including around ~7 dimer repeats along the helical axis (Fig. 2b). Initial Tilt and Psi angle priors are assigned by RELION, which can be used as the basis of later restrained angular searches: 90 ° Tilt from the fact that MTs lie more or less flat on the grid - and Psi from the directionality of helical manual picking coordinates.

As low resolution information is generally sufficient for rough alignment and classification of MTs, particles can usually be binned (we typically bin x 4) until the final fine refinement steps to reduce memory usage and accelerate processing times. Typical pixel size choices of 0.75-1.5Å/pixel result in computationally expensive unbinned box sizes of ~400-800 pixels. To increase signal for classification steps, we generate an additional averaged image for each MT segment by combining 7 neighbouring particles along the helical axis (Fig. 2c). Such segment averages or ‘super-particles’ were already implemented in previous MT processing pipelines, albeit with somewhat varying approaches to their calculation (Sindelar and Downing, 2007; Zhang and Nogales, 2015).

### PF number sorting

A major cause of heterogeneity in an MT dataset is the variable PF number of *in vitro* polymerised MTs. A method for sorting MTs with different PF numbers in 3D was previously developed in Ken Downing’s group, where 6 common MT architectures were described: 11-3, 12-3, 13-3, 14-3, 15-4, and 16-4 (Sui and Downing, 2010), with the first number denoting the PF number and the second the helical start number. Those with an odd start number are pseudo-helical MTs with a seam.

We use a supervised 3D classification approach, similar to that used previously (Sui and Downing, 2010; Zhang and Nogales, 2015), to separate different PF architectures in our cryo-EM data (Fig. 3). To do this, we generate simulated references (see top row, Fig. 3d and Methods) for different PF number MTs (11-16), and compare our experimental MT segment averages to reference projections. Every MT particle is thereby assigned a PF number class. The expectation is that the PF class assignment is the same for all particles from a given MT, but this is not always the case in practice (Fig. 3a). This can be because of the poor signal-to-noise ratio of cryo-EM images, the presence of contamination, distorted or defect-containing MTs not previously excluded during picking, or because of genuine switches in PF number in a single MT. Nevertheless, there is usually a clear dominant PF number class for each MT, and this is imposed on all particles in that MT. We can calculate the confidence we have in each MT PF class assignment by determining the percentage of particles from an MT that fall into the modal PF class (e.g. MTs 1-4 in Fig. 3a have a confidence of 90, 63, 83, and 63 %). By plotting a histogram of the confidence for all MTs in a dataset, it is clear that most PF architectures are assigned with high confidence (Fig. 3b). The use of segment averages increases the confidence in PF number assignment, with 79 % of MTs in the CKK-MT dataset having 100 % confidence when using segment averages versus 71 % using raw MT particles.

**Figure 3.**
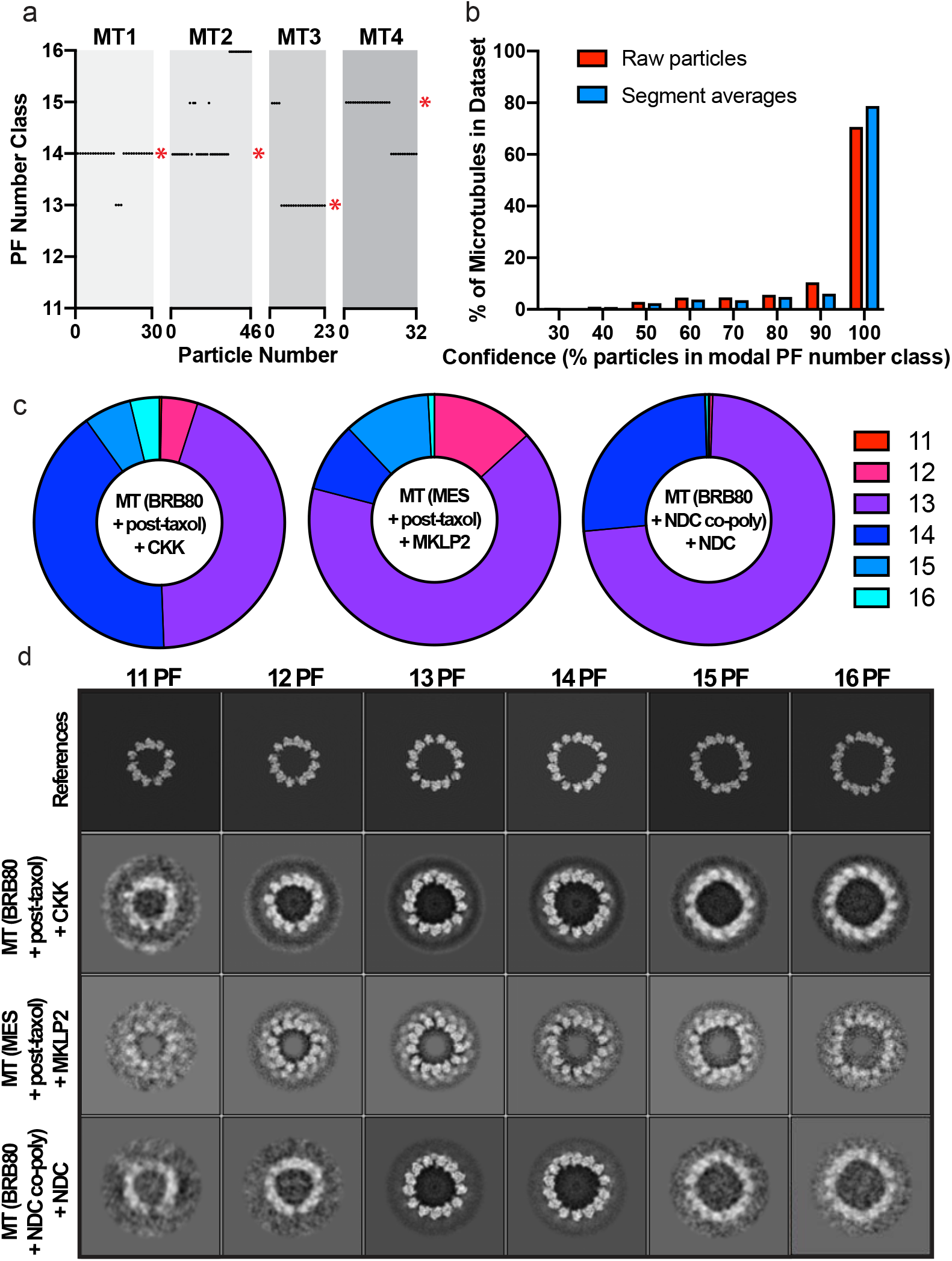
Sorting MTs in 3D by PF architecture. a) Examples of four MTs (MT1-4) from the CKK dataset, showing the PF number class assignment as a function of the particle number within each MT. * indicates the modal class for the microtubule. b) Histogram showing the overall confidence of the MT architecture assignment step (performed with segment averages, or standard particles), plotting the % of MTs within certain confidence values (for the CKK dataset). c) The distributions for 11-3, 12-3, 13-3, 14-3, 15-4, 16-4 PF MTs (% of particles with a certain PF architecture) calculated after MT PF number assignment. d) Central z-axis slices of the different PF number references used for PF number classification, and of the resulting reconstructions for different datasets.

Our three test datasets were analysed using this supervised 3D classification procedure to illustrate that differences in *in vitro* polymerisation methods, which give rise to different distributions of MT PF numbers, are distinguished computationally (Fig. 3c). For the CKK-MT dataset, MTs were polymerised in standard BRB80 buffer then paclitaxel-stabilised, and have a roughly even split between 13-3 and 14-3 PF MTs (44 and 41% respectively). For the MKLP2-MT dataset, MTs were polymerised in a MES-based buffer then paclitaxel-stabilised, resulting in the formation of more 13-3 MTs (66%). MT polymerisation was also performed in the presence of an engineered doublecortin (DCX) chimera (see Methods for details). Polymerisation with the DCX chimera also increased the amount of 13-3 MTs (74 %). In all datasets, there was a minority of other PF types, with 11-3 MTs being very rare.

Fig. 3d shows central slices of 3D references used for supervised classification, and of the resulting PF class reconstructions for CKK-MT, MKLP2-MT, and NDC-MT datasets. The reconstructions show MT structure and PF architecture that matches well with the references, although the less common architectures are less well-defined owing to lower particle numbers and poorer angular coverage in Fourier space.

### Global Search and Initial Seam Assignment

In the next steps, the Euler angles are refined to give an initial estimate of the parameters needed to correctly align each pseudo-helical MT to the reference. The key challenge to address is determination of the Rot angle for each MT that will correctly align the seam of the experimental particles with that of the reference. In other words, although the alignment of multiple different registers of PFs between particle and reference are possible, there is only one PF register that will align the seam accurately. Finding this register is however, very error prone. This is because of the structural similarity of α- and β-tubulin, which means that a Rot angle resulting in a PF register that does not align the seam will still produce a high cross-correlation score.

To determine the Rot angle for each MT, we first perform a ‘global search’ step, composed of two sequential single-iteration 3D refinements to the reference. The first aligns to the reference with a wide search and relatively coarse step (1.8 °, Table 2) in order to assign rough Psi and Tilt angles. The second uses tightly restrained Psi and Tilt angle around those set in the previous alignment, a wide search of the Rot angle and X/Y shifts and a finer angular sampling (0.9 °, Table 2). The result is a non-uniform Rot angle distribution, where the calculated Rot angles form clusters of particles aligned to different PF registers (Fig. 4a). There is a clear bias towards a certain range of Rot angles because PF registers between particle and reference that are closer to aligning the seam will produce higher cross-correlation scores. To give a better initial estimate of the seam location for each MT, we calculate the most commonly assigned Rot angle from the global search step for each MT, and impose it on all particles in that MT (Fig. 4b). In practice, this is still an approximation (as assessed by the suboptimal quality of density of decorating proteins close to the seam in reconstructions at this stage), but assigning a single Rot angle for each MT is essential for later refinement of the seam alignment.

**Figure 4.**
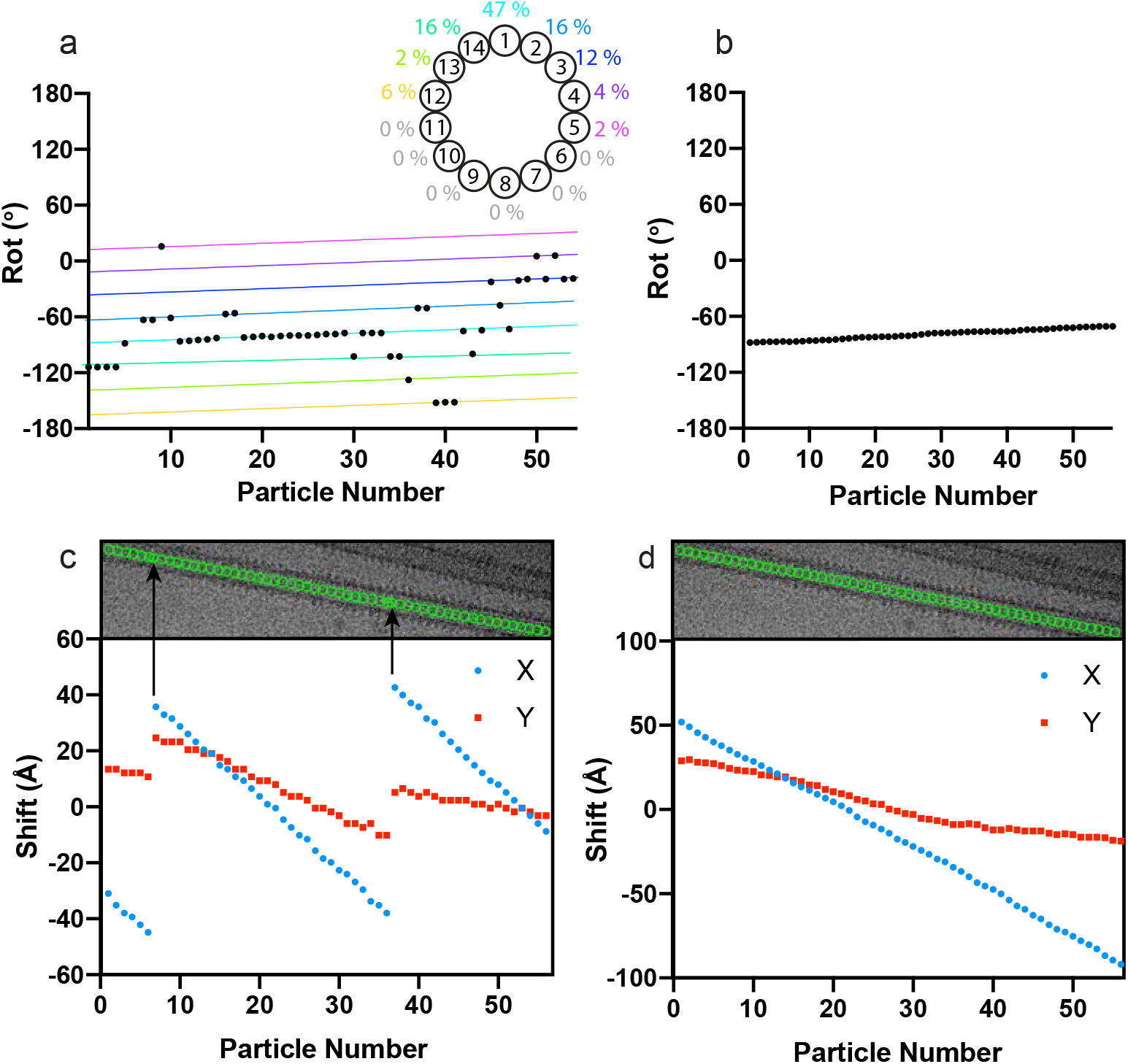
Initial Seam Assignment and X/Y Shift Smoothing. a) MT-Rot angle assignment for a single MT from the CKK dataset, after the ‘Global Search’ step, with the MT Rot angle plotted as a function of the particle number within that MT. The rainbow coloured lines show the clusters calculated during the MT-Rot angle assignment step, with each cluster representing a different PF register being aligned between individual MT particles and the reference – as can be seen by the regular spacing between the clusters. A MT top view representation is annotated by the percentage of particles for this MT aligned with different PFs in the 3D reference. b) Rot angles from the MT example in a), after MT Rot angle assignment. c) The X/Y shifts from the example MT in a-b plotted as a function of particle number. The micrograph of the MT is shown, with the X/Y alignment for each particle in the MT represented by green circles. Particles which have shifted in register are shown by black arrows. d) The X and Y shifts for the example MT in a-c, after X/Y shift smoothing.

### X-Y Shift Smoothing

As described in the pre-processing step, each adjacent particle along a given MT is separated from its neighbours by 82 Å (the approximate length of a α/β-tubulin dimer). The X/Y shifts function to centre each experimental particle with respect to the 3D reference, whilst ensuring that each particle is still approximately 82 Å apart from its neighbours. When plotting X/Y shifts as a function of particle number for a given MT, a single sloping straight line would be expected. What is often observed however, is a large jump in the X/Y shift values (Fig. 4c). These large jumps result in particles being translated along the axis of the MT, causing them to be out of register, either by a tubulin monomer (which can occur because of the similarity of α/β-tubulin) or a tubulin dimer (which will duplicate a neighbouring particle). This occurs even in the presence of MT binding proteins.

To obtain correct X/Y shifts, we first perform a further refinement of X/Y shifts after the initial seam assignment step. Because the wide Rot angle distribution from the global search step also causes a wide X/Y shift distribution, we reset the X/Y shifts to zero, and perform this third alignment, where the Rot/Tilt/Psi angles are locally sampled, but with a wide search range for the X/Y shifts (Table 2). The wide search range is used because manual picking can result in particles which are poorly centred, although this also increases the possibility of mis-translated particles. After this step, mis-translation of particles such as in Fig. 4c is observed. To solve this, we enforce all X/Y shift values in an MT to follow the same slope and intercept, and then perform another refinement (the fourth) with a reduced X/Y shift search area. This ensures every particle is separated by approximately 82 Å (Fig. 4d).

### Refined segment average generation

Once a consistent angle is assigned for all particles along each MT and X/Y shifts are smoothed, a local refinement centred on these parameters is performed using raw particles to ensure all segments are optimally aligned within their local restraints. To maximise signal-to-noise for the subsequent seam check step, new 4 x binned MT segments are extracted centred on these refined translational parameters and new refined segment averages are generated as previously.

### Seam check

To optimise seam allocation for each MT, we took a supervised 3D classification approach using 3D references representing all possible seam positions and their 41 Å shifted positions along the helical axis. The number of references required for a given PF architecture therefore is the number of PFs multiplied by 2, thereby accounting for all possible seam positions and α/β registers. This supervised classification is performed without alignment because we already have crude alignment parameters with internally consistent MT-Rot angles for each MT, and at this stage are further assessing their accuracy. As no alignment is performed, using simulated references of decorating protein alone is beneficial because it represents the most distinctive signal in similar views of MTs rotated around or translated along their helical axis. An example of this approach for the 13PF HsCKK dataset is given in Fig. 5a, showing the CKK-only 3D class references and output reconstructions from all dataset particles classifying into only the most populated 5 classes (Fig. 5a, below).

**Figure 5.**
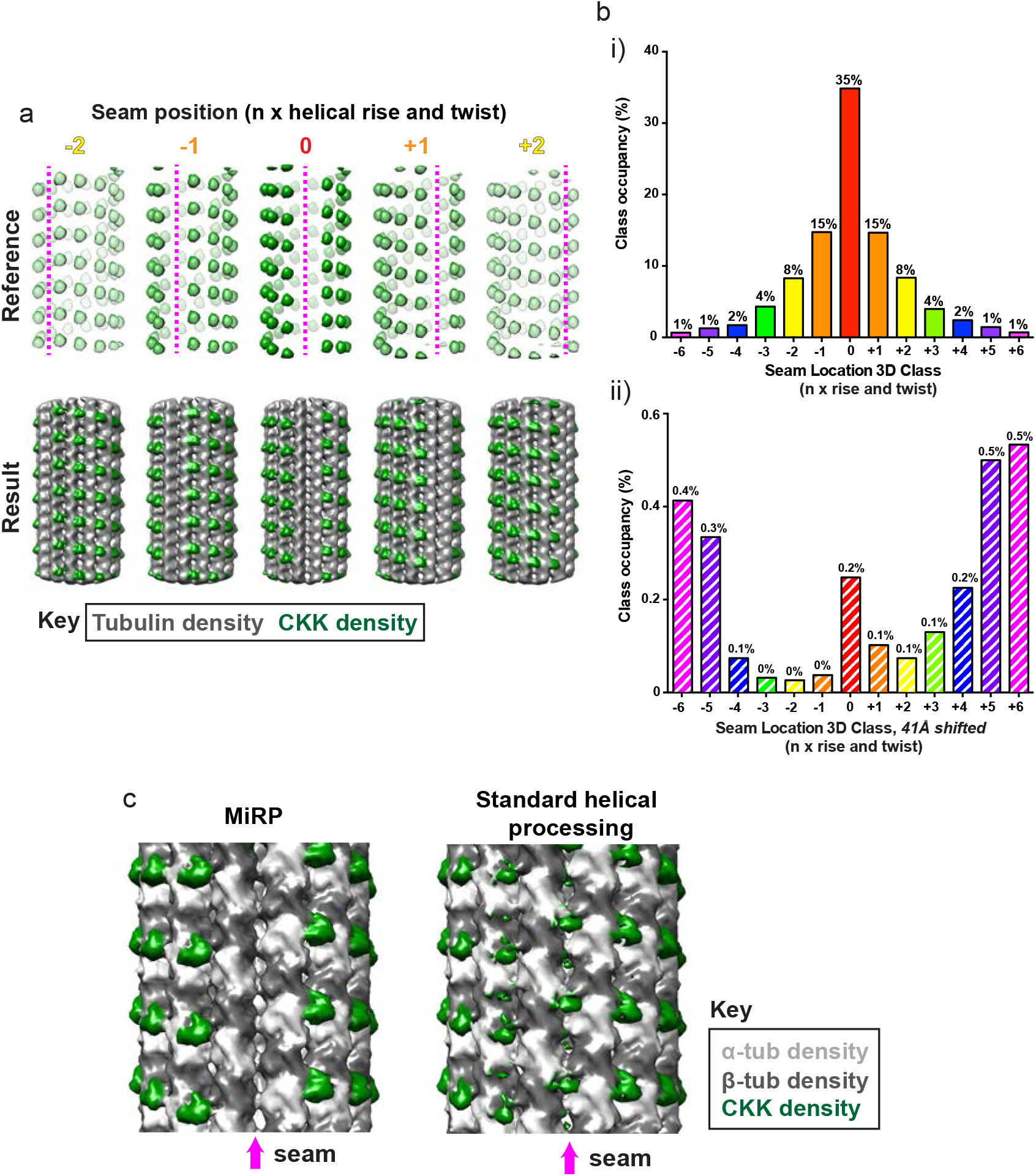
Seam check via supervised 3D classification. a) Supervised 3D classification strategy. Example simulated 13PF CKK density references are shown with rotations around and translations along the helical axis of −2, −1, 0, +1 and +2 times the helical twist and rise. The resulting class reconstructions from a single classification iteration to these references are shown. b) Class occupancy for all 26 supervised classes used for the 13PF CKK-MT dataset. i) Shows classes representing rotations around and translations along the helical axis (angle) of −6 to +6, whereas ii) shows classes with the same rotations and translations plus an additional translation of the monomer repeat distance (41Å). For clarity, class occupancies as a % are indicated above bars representing classes. c) 4 x binned reconstructions of CKK-MT data after application of MiRP and its seam finding procedure or instead using standard helical processing in RELION without intervention. When using MiRP, CKK density is clearly absent from the seam and from 41Å translated locations along the helical axis, whilst aberrant density is found at the seam and 41Å translated locations when using standard helical processing, indicating poor MT Rot angle and αβ-tubulin register determination.

As expected, the most populated class represents correctly aligned data correlating best with the original reference from the previous global search and initial seam assignment steps (Fig. 4). The two next most populated classes represent particles best correlating with this reference rotated and translated either −1 or +1 x the helical rise and twist. This is because the references for these classes are the most similar to the original reference, and thus produce the most likely errors in MT Rot angle assignment. The next two most populated classes include particles best correlating with the original reference rotated and translated either −2 or +2 x the helical rise and twist. This pattern of reducing class occupancy with references rotated and translated increasingly away from the position of the original reference continues, while references shifted a full 41Å along the helical axis are poorly occupied (Fig. 6aii, see scale on the Y axis). The resulting class reconstructions are thus only of good quality in the best occupied classes, and the poorly occupied classes contain too few particles for either good signal-to-noise or angular distribution.

**Figure 6.**
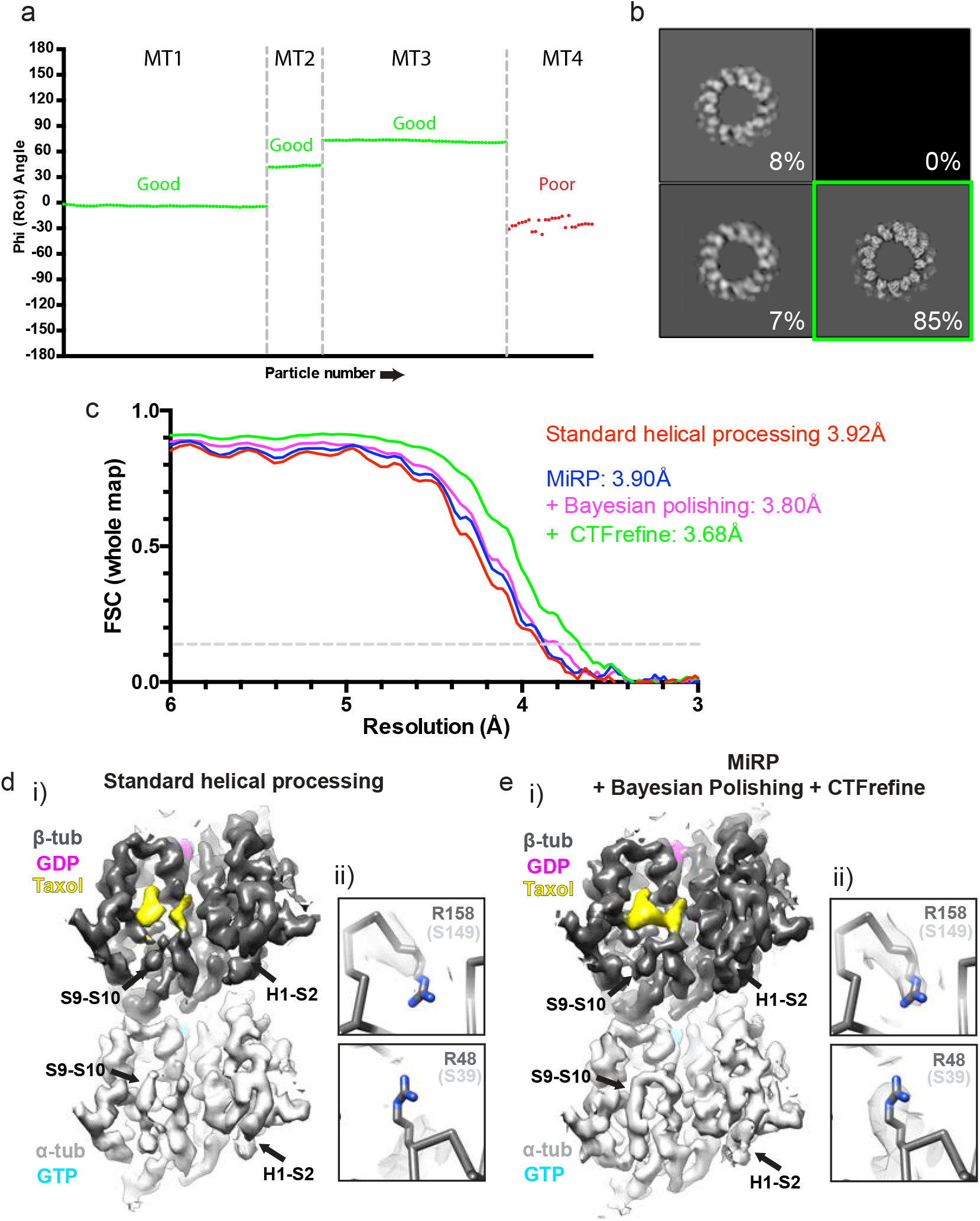
Data optimisation and high-resolution reconstruction. a) Plot showing particle angles for 4 13PF MTs from the MKLP2-MT dataset as a function of particle number (neighbouring particles are separated by ~82Å along the helical axis). Three MTs have aligned as expected (‘good’) whilst one shows jumps in angle along the helical axis and can be excluded (‘bad’). b) Result of the 3D classification without alignment to a preliminary unbinned reconstruction reference of 13PF MKLP2-MTs, displayed as 2D slices in the RELION display GUI. 85% of particles went into a ‘good’ class with expected structural features and a defined seam (green box) and were taken for further processing. c) Gold-standard corrected FSC curves for the 13PF CKK-MT dataset from RELION post-processing, using the central masked 15 % of the reconstructions along the helical axis (~90Å, a little over the dimer repeat distance). Final symmetrised reconstructions are compared after standard helical processing (without Bayesian polishing or CTF refinement) or after use of MiRP without or with Bayesian polishing or Bayesian polishing and CTF refinement. d) Unique features of α- and β-tubulin are poorly resolved after standard helical processing in RELION. i) The lumenal face of the tubulin dimer of the asymmetric unit opposite the seam for symmetrised 13PF CKK-MT reconstructions showing poorly defined density for the H1-S2 and S9-S10 loops, which are distinct in α and β-tubulin. ii) Density for non-conserved α and β-tubulin sidechains such as β-tubulin’s R158 (S149 in α-tubulin) and R48 (S39 in α-tubulin) are poorly defined. e) Unique features to α and β-tubulin are well resolved after application of MiRP. i) The lumenal face of the tubulin dimer of the asymmetric unit opposite the seam for symmetrised 13PF CKK-MT reconstructions exhibits well defined density for the H1-S2 and S9-S10 loops, which are distinct in α- and β-tubulin. ii) Density for non-conserved α- and β-tubulin sidechains such as β-tubulin’s R158 (S149 in α-tubulin) and R48 (S39 in α-tubulin) are well defined.

Some samples produce sharper peaks around the original ‘central’ reference than others, reflecting the success of the initial MT-Rot angle assignment in the initial seam identification step. In the three test datasets used here, there is a rough correlation between increasing size of the regularly decorating part of the protein (estimated molecular weight of ordered region in reconstructions: N-DC 10 kDa, CKK 14 kDa, MKLP2 38 kDa) and the sharpness and height of the alignment peak - this would be expected from the increased protein signal used to discriminate the correct Rot angle (Supplementary Fig. 1). However, other factors like decorating protein occupancy, ice-thickness data collection parameters and detector quantum efficiency (DQE) are likely to play a role.

As the input MT-Rot angles are unified for each MT, and seam position changes are likely to be very rare along MTs in standard *in vitro* preparations (Chretien and Fuller, 2000), we also unify class allocations determined by the seam finding step for each MT. The class allocations for each MT segment in a given MT are adjusted to the mode for all segments in that MT. According to the unified class allocation for each MT, which indicate deviation from the true seam position and α/β register, particle angles and translations along the helical axis are adjusted in the output parameter file (.star file) using multiples of the helical twist and rise, similar to the correction strategy applied by Zhang & Nogales (Zhang and Nogales, 2015). The lack of reference bias in classification and success of these angular and translational adjustments are illustrated in multi-iteration local C1 3D refinements with particles from a single unified class to references with a new seam position (Supplementary Fig 2). In these cases, the seam location in the resulting reconstructions, remains at the position of the class reference from the 3D classification step despite the new reference having a different seam position. In contrast, the Rot angle and translational correction adjusts the seam position in the output reconstructions as expected, even when the new reference has a different seam position. The output reconstructions exhibit subnanometer resolution features when compared to the 15 Å low-pass filtered references, further reinforcing the lack of reference bias. The single 3D classification iteration used at the seam check step thus gives reliable class assignments and avoids poor quality anisotropic references generated in poorly occupied classes being used in further iterations.

At this stage, the success of the protocol can be assessed in the quality of output reconstructions. A crude reconstruction using the raw particles of the binned data should give a clearly identifiable ‘clean’ seam indicating low error in angle assignment. Using the protocol with a dataset of 13PF MTs decorated with the CAMSAP CKK gave a clear pattern of CKK binding every 82Å between tubulin dimers and an absence of CKK density at the seam or at the structurally equivalent binding site 41Å along the helical axis (particularly at or close to the seam, Fig. 5c, left). In contrast, standard helical processing in RELION was less successful as indicated by incorrect CKK density at the predicted seam position and at 41Å shifted positions along the helical axis (Fig. 5c, right). These incorrect densities arise from MT Rot angle and α/β register assignment errors, while their absence when using MiRP supports our hypothesis that this process improves MT Rot angle and α/β register assignment.

### Data optimisation and high-resolution reconstruction

After determination of rough alignment parameters using binned data as described, the data are cleaned to exclude putative poor particles (such as misaligned, contaminated or low-resolution particles) from the final dataset for high-resolution reconstruction. Firstly, alignment quality within each MT is checked against our expectation that neighbouring MT segments have similar angular and translational assignments. For example, plotting particle number against Rot angle for each MT (Fig. 6a) helps identify MTs with poorly aligned segments by observing angles that deviate significantly between neighbouring MT segments. These MTs are removed.

A final dataset containing selected good MTs are subjected to high-resolution C1 refinement using recentered (according to previously determined translations) unbinned data and RELION’s auto-refinement procedure with tightly restrained rotation and translational searches around the previously determined parameters. At this stage, a 3D classification without alignment (~25 iterations or as many as required for convergence) can also be run to allow poor or low-resolution 3D classes to be removed (Fig. 6b). The remaining high-resolution classes are then subjected to a final high-resolution asymmetric refinement or a refinement with the appropriate helical symmetry applied. An asymmetric refinement is recommended first, as the C1 reconstruction represents the true MT structure and can be used to determine approximate helical parameters for symmetrisation. Furthermore, the quality of the seam in the C1 reconstruction is a good indication of the success of the applied MiRP procedure. Identification at this point of appropriate initial helical parameters and local search ranges for helical parameter refinement in RELION is vital for final refinements with applied helical symmetry.

During a successful final asymmetric refinement, reconstructions with a ‘clean’ seam should be generated during each auto-refine iteration (used as references for the next iteration). In contrast, due to the pseudo-symmetric nature of MTs with a seam, reconstructions calculated with applied helical symmetry during refinement will only have good density for unique α and β tubulin features and decorating proteins (if binding every dimer) in the region of the reconstruction opposite the original seam position. This is because opposite the seam is the only position in the MT where no averaging of non-identical subunits across the helical symmetry break at the seam occurs. Conversely, all tubulin subunits in all PFs retain their fundamental, conserved tubulin 3D structure. The use of symmetrised rather than pseudo-symmetrised references (as in previous MT processing pipelines, (Sindelar and Downing, 2007; Zhang and Nogales, 2015)) doesn’t matter in practice, as translational and rotational searches are highly restrained during this final local refinement. As we are only interested in the structure of the asymmetric unit in symmetrised reconstructions, the ‘good’ asymmetric unit opposite the original seam position can be extracted and used for structural analysis.

In order to improve the resolution of the final reconstructions, RELION v3.0’s Bayesian polishing and CTF refinement modules (Zivanov et al., 2018) are used. MTs are large objects that provide good signal for these per-particle approaches, allowing good local determination and correction of particle motions and CTF parameters. In our hands, with near-atomic resolution data, a single iteration of Bayesian polishing followed by CTF refinement followed by 3D auto refinement, applied post-pipeline, results in minor improvements to resolution (~0.2Å), with the CTF refinement step being the most effective (Fig. 6c). Whilst measured resolution improvement is minor, Bayesian Polishing and CTF refinement can improve visualization of key structural features (Supplementary Fig. 3).

Strikingly, although application of MiRP compared to standard helical processing has a negligible effect on the reported reconstruction resolution by FSC (Fig. 6c), the structural details are clearly superior in quality. Whilst the expected density features unique to α- or β- tubulin in the symmetrised ‘good’ asymmetric unit opposite the seam are poorly resolved and indistinct when using standard helical processing in RELION (Fig. 6d), they are well resolved and distinct with application of MiRP (Fig. 6e). For example, successful application of the protocol resulted in improved definition of the S9-S10 and H1-S2 loops – which differ structurally in α and β tubulin - and bound paclitaxel (binds specifically to β-tubulin) in symmetrised reconstructions of 13PF CAMSAP-CKK decorated MTs, when compared to using the standard helical processing (Fig. 6di,ei). Furthermore, specific side chains that are different between α- and β-tubulin are better resolved (Fig. 5dii,eii). The improvement in local map to model cross-correlation in the decorating protein and in the structurally distinctive α and β tubulin features is further evidence of successful data alignment by MiRP compared to standard helical processing (Supplementary Fig. 4).

### Application to 3 test datasets

C1 reconstructions of all test datasets showed that the MT seam was well determined, independently of whether the decorating protein binds within the inter-PF groove (Fig. 7a,c) or on the PF ridge (Fig. 7b). The asymmetric units of symmetrised reconstructions gave near-atomic resolution detail in all datasets (Table 4), with β-sheet separation and helical pitch clearly resolved (Fig. 7d,e,f). A consistent feature in all reconstructions is a resolution gradient where tubulin is best resolved (particularly in the core) and the decorating protein is less well resolved, particularly at distal regions (Fig. 7g,h,i).

**Figure 7.**
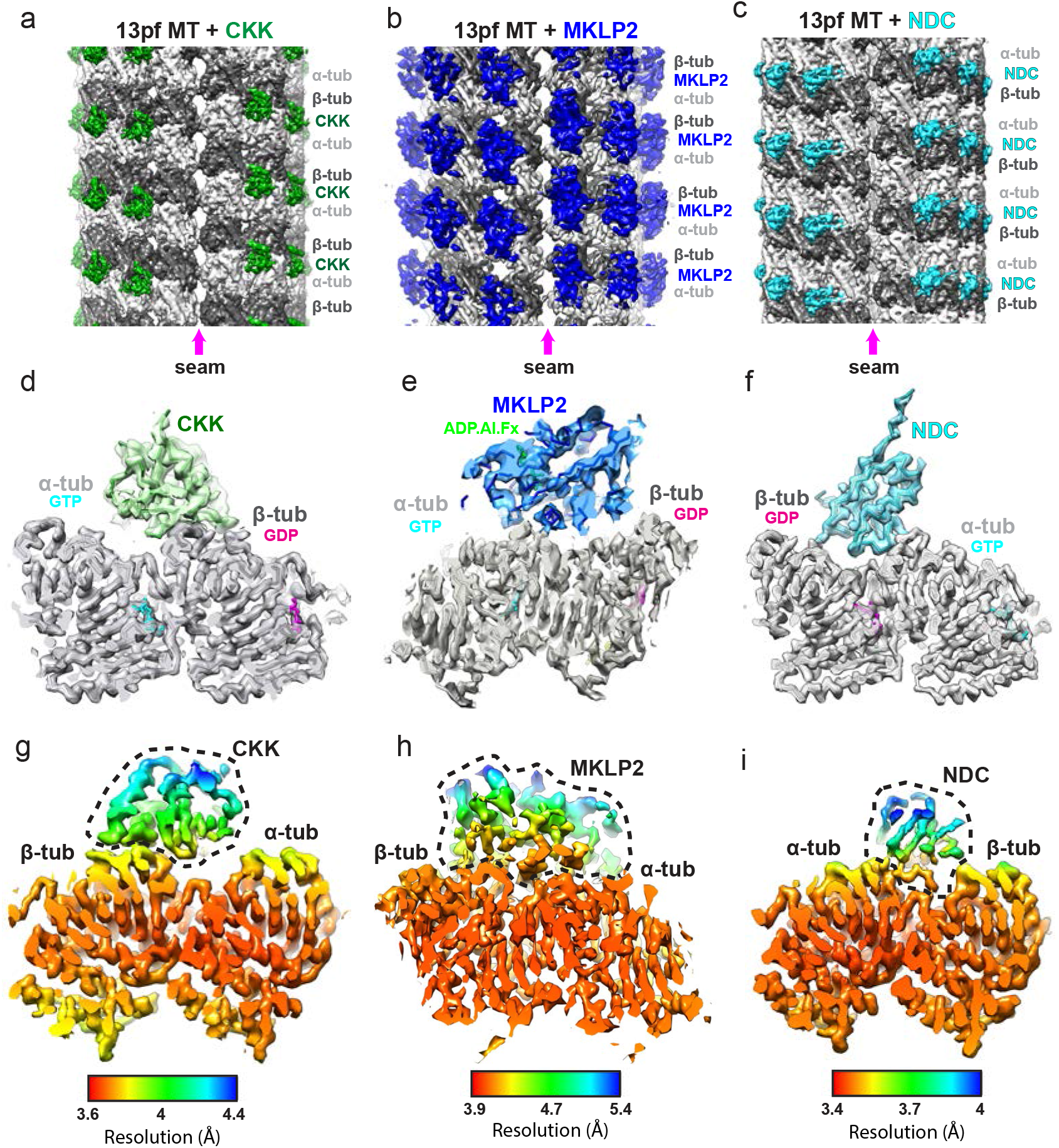
Final reconstruction results for test datasets. a) C1 reconstruction of the 13PF CKK-MT dataset (unfiltered), showing a well-defined seam indicative of accurate MT Rot angle and αβ-tubulin register assignment. b) C1 reconstruction of the 13PF MKLP2-MT dataset (unfiltered), showing a well-defined seam indicative of accurate MT Rot angle and αβ-tubulin register assignment. c) C1 reconstruction of the 13PF NDC-MT dataset (unfiltered), showing a well-defined seam indicative of accurate MT Rot angle and αβ-tubulin register assignment. d) Density and fitted model for the ‘good’ asymmetric unit opposite the seam in the symmetrised reconstruction of the 13PF CKK-MT dataset, showing density quality consistent with the reported resolution. e) Density and fitted model for the ‘good’ asymmetric unit opposite the seam in the symmetrised reconstruction of the 13PF MKLP2-MT dataset, showing density quality consistent with the reported resolution. f) Density and fitted model for the ‘good’ asymmetric unit opposite the seam in the symmetrised reconstruction of the 13PF NDC-MT dataset, showing density quality consistent with the reported resolution. g) As in panel e but from a different viewpoint, showing local resolution determined by RELION’s local resolution software. The CKK decorating protein is within the dashed black line. h) As in panel e, but from a different viewpoint showing local resolution determined by RELION’s local resolution software. The MKLP2 decorating protein is within the dashed black line. i) As in panel f, but from a different viewpoint showing local resolution determined by RELION’s local resolution software. The NDC decorating protein is within the dashed black line.

**Table 4.** Dataset size and resolutions. Gold-standard corrected FSC resolution at the 0.143 threshold for symmetrised reconstructions calculated with RELION post-processing, using the central masked 15 % of the reconstructions along the helical axis (~90Å, a little over the dimer repeat distance).

| Decorator | Number of selected micrographs | Starting particle number | Final 13pf particle number | Final 13pf resolution (Gold standard FSC 0.143) |
| --- | --- | --- | --- | --- |
| CKK domain of CAMSAP1 | 1075 | 82,666 | 26,854 | 3.68Å |
| Motor domain of MKLP2 | 293 | 25,568 | 14,411 | 4.19Å |
| N-DC domain of DCX | 847 | 50,255 | 28,347 | 3.63Å |

## Discussion

Cryo-EM has been a critical methodology for revealing detailed mechanisms of MT dynamic instability (Alushin et al., 2014; Chaaban et al., 2018; Manka and Moores, 2018a; Manka and Moores, 2018b; Zhang et al., 2015), characterising the interactions of MT-associated proteins important for a plethora of different biological processes (Amos and Schlieper, 2005; Downing and Nogales, 2010; Manka and Moores, 2018a; Nogales and Kellogg, 2017; Nogales and Zhang, 2016), and determining the mechanism of important MT-binding drugs such as paclitaxel (Alushin et al., 2014; Kellogg et al., 2017). Whilst true helices give many identical views around their axis and have been used to determine near-atomic resolution MT structures (Benoit et al., 2018), the majority of MTs polymerised *in vitro* have lattice architecture with a symmetry break at a discontinuity known as the seam. This results in many highly similar but unique views that are difficult to discriminate from one another in the noisy 2D projection cryo-EM images, and which presents a specific challenge in structure determination.

Initial approaches to determining the structure of undecorated pseudo-helical MTs treated them as if they were helical, resulting in the loss of distinctive α and β tubulin density and of a defined seam in the reconstructions (Li et al., 2002; Sui and Downing, 2010)). Use of a fiducial protein binding every tubulin dimer facilitates identification of the seam position, but inaccuracies in α and β tubulin register and MT-Rot angle assignment remain, causing poor seam definition and α/β-tubulin differentiation. As shown previously (Zhang and Nogales, 2015) and in our current work, this loss of reconstruction quality is not particularly reflected by resolution loss as judged by FSC. To our knowledge, two dedicated pipelines for processing pseudo-helical MTs have been developed: by Charles Sindelar and Ken Downing (Sindelar and Downing, 2007) for use with SPIDER (Frank et al., 1996) and Frealign (Grigorieff, 2007), and by the Nogales group (Alushin et al., 2014; Zhang et al., 2015; Zhang and Nogales, 2015) for use with EMAN (Ludtke et al., 1999) and Frealign. The first pipeline has been used to solve a number of pseudo-helical MT structures with fiducial decorating proteins (Shang et al., 2014; Sindelar and Downing, 2007; Sindelar and Downing, 2010) - including by our group e.g. (Atherton et al., 2014; Atherton et al., 2017a; Fourniol et al., 2010; Goulet et al., 2012; Manka and Moores, 2018b; Maurer et al., 2012) - to subnanometer or near-atomic resolutions. The Nogales group pipeline has been used to solve pseudo-helical MT structures with or without fiducial decorating proteins to near-atomic resolutions e.g. (Alushin et al., 2014; Kellogg et al., 2017; Kellogg et al., 2018; Kellogg et al., 2016; Zhang et al., 2015; Zhang et al., 2018; Zhang et al., 2017).

RELION is very popular software for cryo-EM image processing (Patwardhan, 2017) and when helical processing options were added, we became interested in using it to process MTs. Movie alignment and CTF determination are built-in modules in RELION and are compatible with later operations performed on individual particles, requiring no scripting and little intervention from the user. RELION is also a GPU enabled program, vastly reducing processing times for computationally expensive MiRP steps. Building on previously established principles in other pipelines, we have developed a pipeline for processing MTs completely within RELION, with scripted interventions on parameter files and particle images and imposing control over advanced options for restrained alignments.

In MiRP, MT segments picked manually in RELION are extracted, adjacent segments averaged to improve signal-to-noise (similar to previous approaches (Sindelar and Downing, 2007; Zhang and Nogales, 2015)), then supervised 3D classification performed to identify MT PF number architecture (similar to previous approaches (Bell et al., 2017; Sui and Downing, 2010; Zhang and Nogales, 2015)). Subsequently, to deal with the major challenge of accurately aligning and reconstructing the seam, we perform a global search of alignment parameters, followed by optimisation of MT Rot angle and αβ-tubulin register assignment, then an explicit seam checking step. Both the Sindelar and Nogales group pipelines perform similar steps, albeit with different implementations. The Sindelar pipeline performs seam finding based on the angular assignments in the global search step; in the process, MTs with poor internal angular consistency are excluded, while Euler angles and X/Y shifts in MTs to be included are smoothed by least squares trimming (Liu et al., 2017; Sindelar and Downing, 2007). While we do not routinely exclude MTs in this way, this may be advantageous particularly for big datasets where automated data processing is important, and we have written scripts implementing this step in MiRP. After the global search step, the Nogales pipeline optimises initial seam assignment, and then performs an explicit seam finding step by testing, for each particle against a single reference, parameters representing each possible seam location. This approach inspired ours, where we instead perform seam finding based on supervised 3D classification of a single set of particle parameters against multiple references representing all possible seam locations.

Once rough alignment parameters are determined with binned data, we perform final refinements with RELION’s auto-refine procedure. Local refinement restrained around rough parameters can be performed much like in Frealign used in previous pipelines; however the RELION procedure requires less user intervention. RELION’s helical mode also has the advantage of iteratively refining helical parameters during the refinement procedure internally, whereas previous pipelines required helical parameter determination and iterative refinement to be outsourced. Furthermore, RELION v3.0 implements CTF refinement and Bayesian polishing modules that, in our hands, resulted in further - albeit minor - reconstruction improvements.

One key difference between the current pipeline and previous ones is that we do not apply pseudo-symmetry to generate new references every iteration during the refinement procedure. Previous approaches applied full helical symmetry to reconstructions generated during the procedure, then reproduce the ‘good’ helical subunit opposite the seam around the helical axis, to generate new references for each iteration. After the final refinement iteration either C1 or pseudo-symmetrised maps are generated (Sindelar and Downing, 2007; Zhang and Nogales, 2015). In the initial stages of our approach, we perform single iterations of rough alignment to a simulated reference, and do not use references derived from the data. During these initial stages, output C1 reconstructions are only used for checking the success of the MiRP procedures. Once rough alignment parameters are determined, only local alignments of the unbinned data are performed, such that deviations of Rot angles and translations more than the helical twist or monomer repeat distance (41Å) are not allowed. During the multi-iteration C1 refinement of unbinned data, new C1 reconstructions with a seam are generated from the data every iteration as references for the next. In terms of retaining the true C1 structure of the MT, this approach is the only way to avoid introducing artefacts by incorrect symmetry allocation, because no symmetry is applied either to the final structure or to references generated during refinement iterations. Such C1 reconstructions can reveal subtle variation within helical parameters between subunits around the helical axis lost after symmetrisation or pseudo-symmetrisation (Zhang et al., 2015; Zhang et al., 2018). For optimal C1 refinements, references should match the experimental projection data as closely as possible - thus references with these variations in helical parameters are retained in our MiRP procedure.

The final C1 reconstruction generated by MiRP can be used to determine average initial helical symmetry parameters for input. During the subsequent multi-iteration refinement that applies symmetry with the objective of improving the quality and resolution of the final asymmetric unit reconstruction, the MiRP user provides rough symmetry parameters, and helical symmetry is applied and refined every iteration within RELION. The larger the variation in helical parameters around the circumference, the less symmetrisation will improve asymmetric subunit density and resolution. Regardless, the application of helical symmetry means that all asymmetric units but the one opposite the seam are distorted by averaging over the seam. In our procedure, symmetrised new references generated every iteration as input for the following iteration are thus not representative of the true structure. However, due to their close structural similarity, the resulting blur of α and β-tubulin subunits nevertheless produces references each iteration able to drive restrained local refinement to near-atomic resolutions. In other pipelines, a simulated pseudo-helical MT is built by copying the single ‘good’ asymmetric unit around the MT circumference using helical parameters. However, as a high-resolution asymmetric unit is the objective of symmetrisation, we reason that copying the ‘good’ asymmetric unit round the MT cirumference using helical parameters is unnecessary. It should also be noted that pseudo-symmetrisation procedures are potentially error prone and can lead to misleading or suboptimal structures if helical parameters and ranges for their refinement are not assigned appropriately.

An alternative approach from the Carter group, implemented whilst this manuscript was in preparation, performs local averaging of user-defined helical subunits in real-space in RELION (Lacey et al., 2019). This could potentially account for helical parameter variation around the MT circumference - it therefore represents a method for generating pseudo-symmetrised MT reconstructions and benefits from improvements from subunit averaging whilst retaining helical parameter variations. The approach was used to determine the structures of dynein fragment-bound pseudo-helical MTs at ~4-5Å without some of the specific interventions implemented in MiRP. This approach was reported by the authors to work well for the dynein datasets but produced sub-optimal seam determination for an EB dataset as judged by seam-quality in C1 reconstructions. Therefore, although this alternative approach requires less user intervention, we expect that, in particular for smaller (<15kD) decorating proteins, data processing in MiRP will improve MT Rot angle calculation and αβ tubulin register determination, and therefore the final reconstruction quality. Nonetheless, the Carter group pseudo-symmetrisation approach may be useful when applied during final high-resolution refinement with symmetry at the high-resolution reconstruction stage of the MiRP procedure. However, caution is required as the reliability and benefits of this process will be highly dependent on how well the subtle variations in helical parameters around the MT are determined and applied.

Applying MiRP to 3 modestly sized MT datasets (10,000-30,000 final particles) manually collected on two different direct electron detectors resulted in near-atomic resolutions reconstructions. The resolution of MKLP2-MT dataset was improved ~0.2Å and the density quality markedly improved over that previously published by our group using the Sindelar pipeline (Atherton et al., 2017b). In addition, the CKK-MT dataset was improved ~0.5Å in resolution and the density, particularly for the decorating proteins, noticeably improved over previous attempts with the Sindelar pipeline (data not shown). In all three datasets processed using MiRP, the seam was well-defined in the C1 reconstructions, and yielded high-quality density for decorating proteins and unique α or β-tubulin features in the resulting symmetrised reconstructions.

The Nogales group protocol has been successfully applied to datasets of undecorated MTs (Zhang et al., 2018). Our current protocol is optimised for processing MTs with fiducial decorating proteins binding every tubulin dimer but, with minor modifications, we have also determined the structure of the MT-bound EML4 N-terminus which binds every tubulin monomer (Adib et al., 2019). Proteins bound every monomer do not act as effective fiducials but rather add noise to the data - they therefore pose a similar or even bigger challenge in comparison to processing undecorated MTs, but our EML4-MT reconstruction points to the power and applicability of MiRP.

RELION is regularly improved and updated, therefore future iterations of MiRP can be adapted to best make use of further improvements in cryo-EM image processing. Furthermore, it should be noted that presently MiRP treats all asymmetric units in MT segments as a single population during classification and refinement. In the future, we aim to harness the power of RELION to expand MiRP to analyse heterogeneity of the asymmetric units within MTs and thereby open up this poorly understood aspect of MT biology.

## Acknowledgements

This work was funded by a grant from the Medical Research Council, U.K. to C.A.M (MR/R000352/1) and a PhD studentship to A.D.C. (MR/J003867/1). EM data collection was supported by grants from the Wellcome Trust (079605/Z/06/Z, 101488/Z/13/Z) and the BBSRC (BB/L014211/1).

---

**Supplementary Figure 1.**
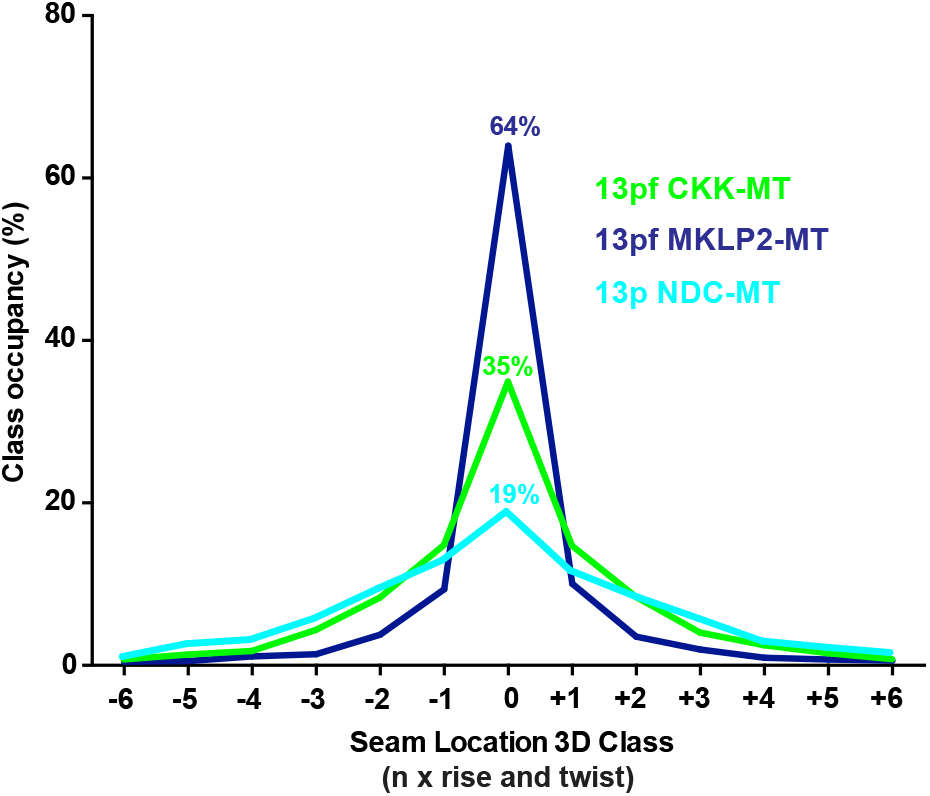
Seam finding 3D class allocation distribution for the 3 test datasets. Class occupancy distribution of 13PF particles from CKK, MKLP2 and NDC decorated datasets classifying to 13PF references built from appropriate decorating protein only density, with seams in modified positions (modified by −6 to +6 multiples of the helical rise and twist).

**Supplementary Figure 2.**
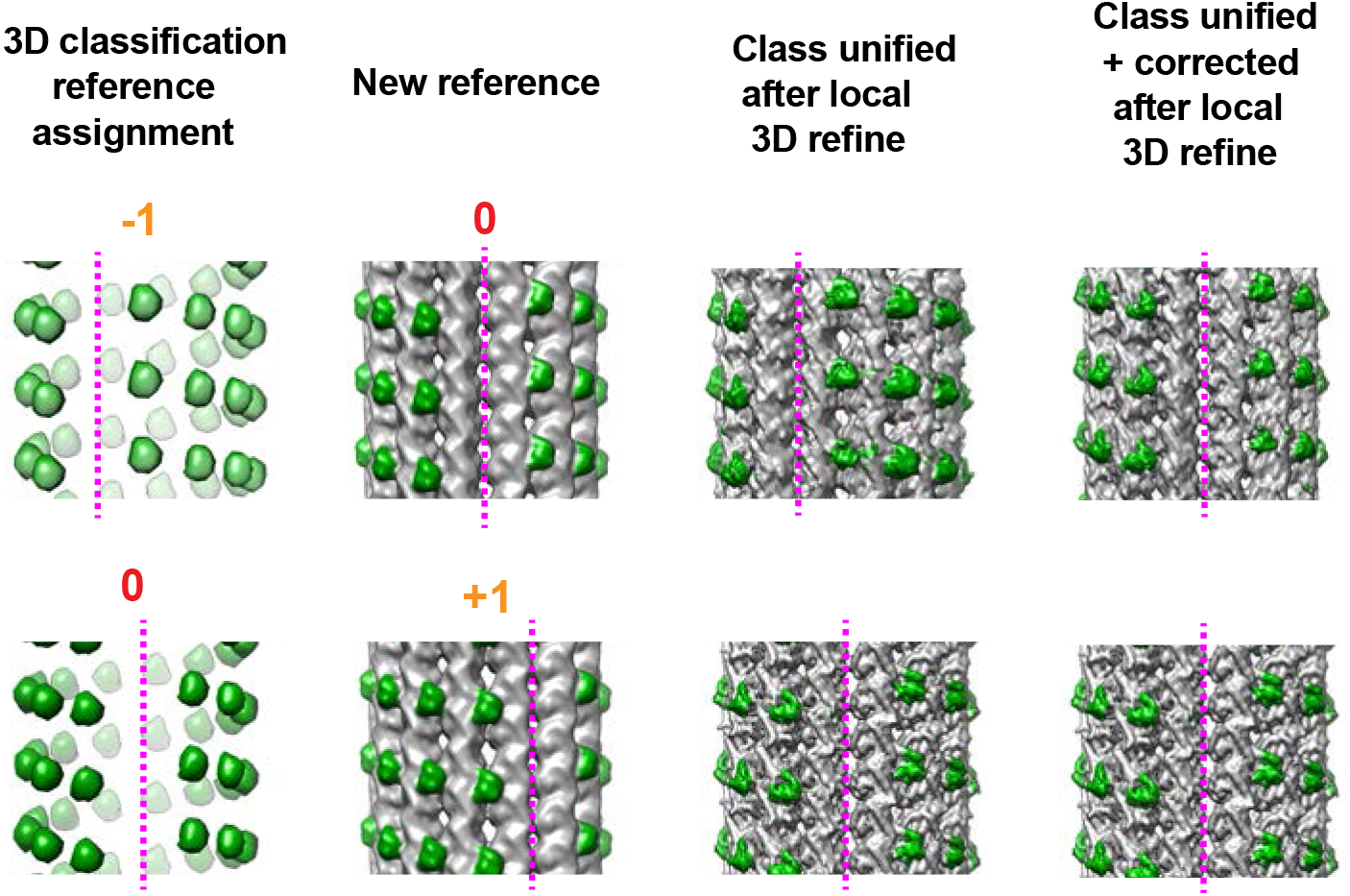
Testing the seam check 3D classification and MT-Rot angle/translational correction procedure. Unbinned datasets corresponding to MTs (particles after per-MT unification), classifying to CKK decorating protein only 3D references, with seams in modified positions (modified by −1 to 0 multiples of the helical rise and twist) shown in the first column were extracted. These datasets were subjected to multi-iteration local C1 refinements to new unbinned references, shown in column 2, of tubulin with decorating protein with a modified seam position. The 3D reconstructions from these refinements without (column 3) or with (column 4) correction of the MT Rot angles and translations are shown.

**Supplementary Figure 3.**
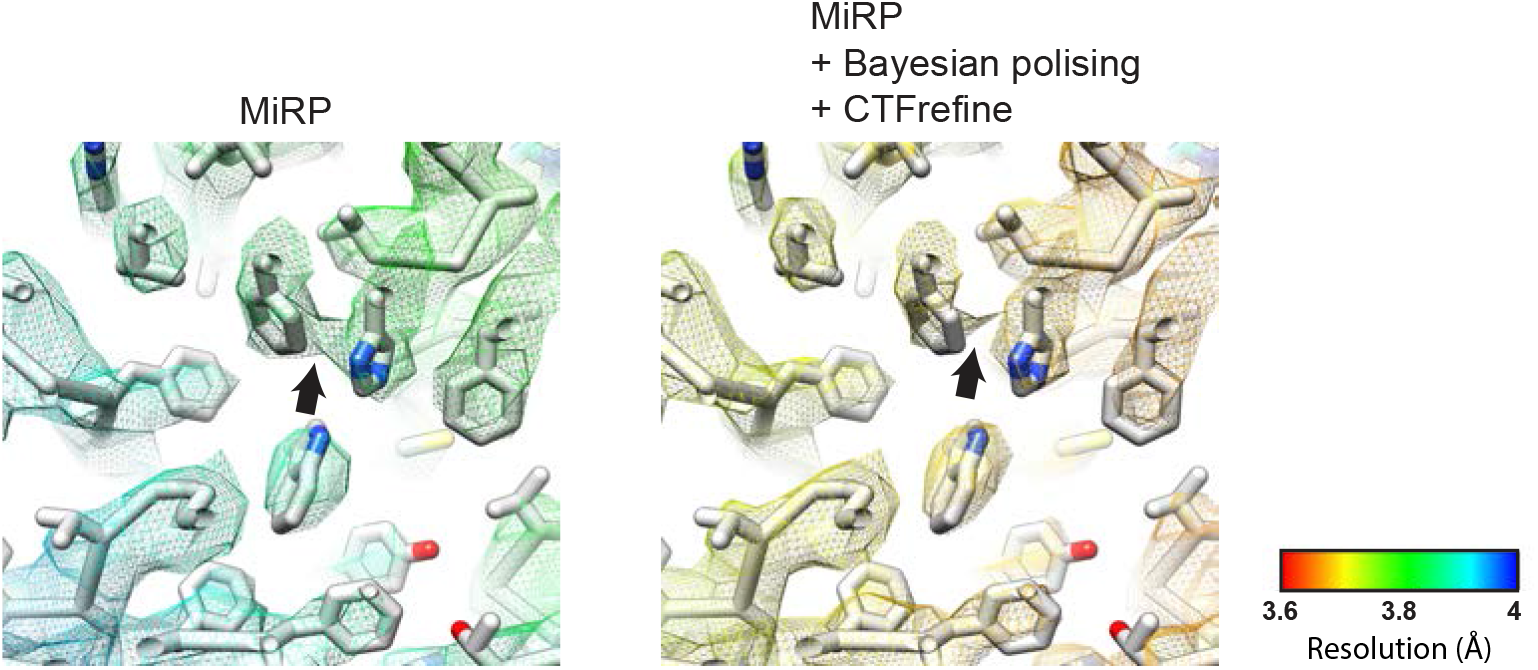
Resolution improvements after Bayesian polishing and CTF refinement. A central portion of α-tubulin model and density from the symmetrised CKK-decorated asymmetric unit (density shown as mesh), coloured by local resolution, is shown upon completion of the MiRP procedure at equivalent thresholds preceding or following single iterations of Bayesian polishing and CTF refinement procedures in Relion.

**Supplementary Figure 4.**
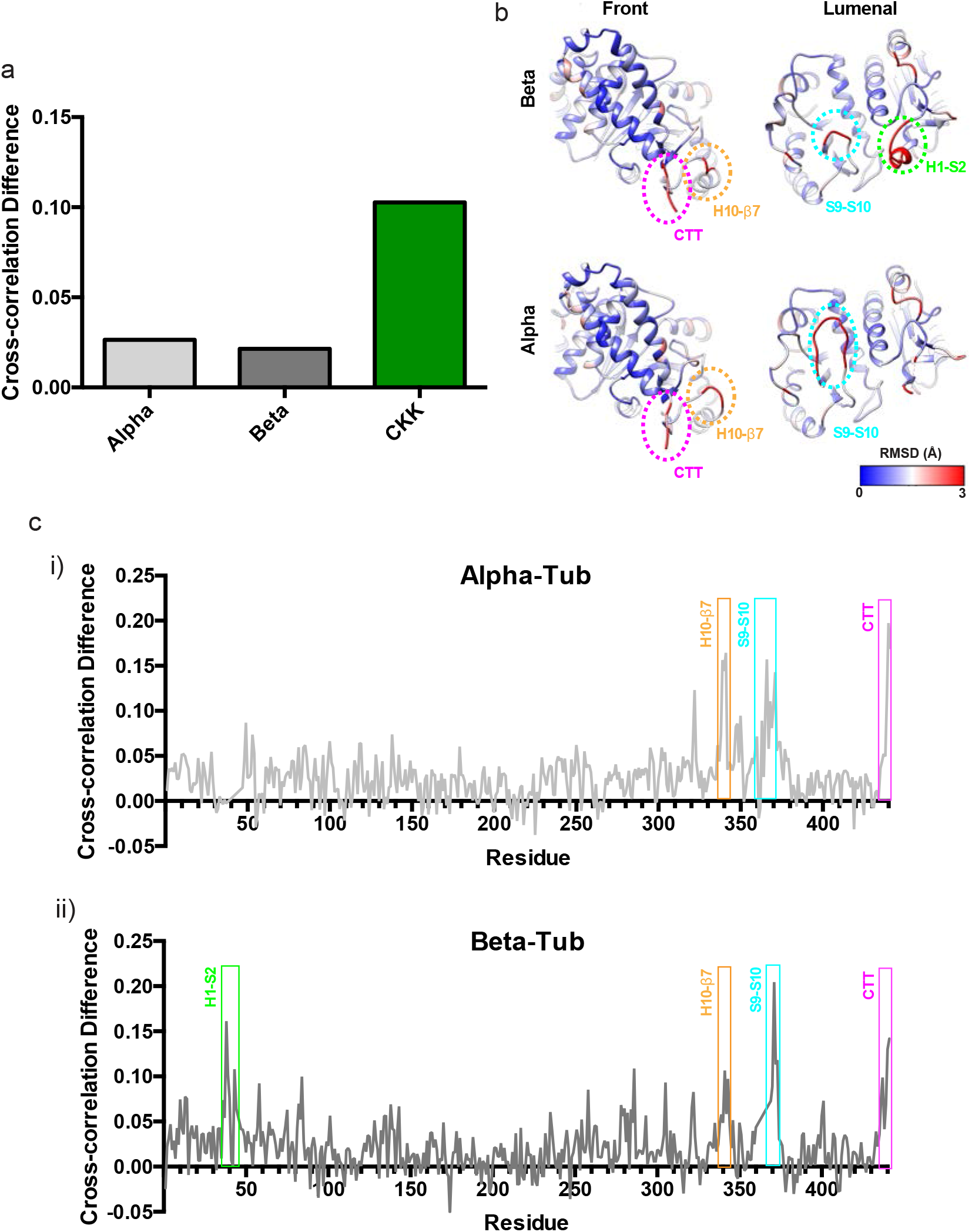
CKK-MT dataset map to model cross-correlations demonstrate that MiRP improves MT-Rot angle and αβ-tubulin register determination. a) Map to model local cross-correlations were calculated between the model built from MiRP-derived density (Atherton et al., 2019) and the CKK-decorated asymmetric unit density derived from either standard helical processing or the MiRP procedure. The former values were then subtracted from the latter to give cross-correlation differences. Asymmetric unit cross-correlation improves most for the CKK region of the reconstruction because it is most sensitive to the successful alignment performed by MiRP; global tubulin density and therefore model fitting is always good, even in reconstructions calculated using standard helical processing. b) Model RMSD between superimposed α and β-tubulin models (MiRP-derived) shown on either front or lumenal faces of α or β-tubulin. Structural regions of particularly high RMSD, indicating significant structural differences, are indicated with dashed rings. c) Per-residue differences in cross-correlation show local improvements from the MiRP procedure for i) α and ii) β-tubulin, with improvements highest for indicated regions of high structural divergence corresponding to those identified in panel b.

